# Population genomic history of the endangered Anatolian and Cyprian mouflons in relation to worldwide wild, feral and domestic sheep lineages

**DOI:** 10.1101/2023.11.23.568468

**Authors:** Gözde Atağ, Damla Kaptan, Eren Yüncü, Kıvılcım Başak Vural, Paolo Mereu, Monica Pirastru, Mario Barbato, Giovanni Giuseppe Leoni, Merve N. Güler, Tuğçe Er, Elifnaz Eker, Tunca Deniz Yazıcı, Muhammed Sıddık Kılıç, N. Ezgi Altınışık, Ecem Ayşe Çelik, Pedro Morell Miranda, Marianne Dehasque, Viviana Floridia, Anders Götherström, C. Can Bilgin, İnci Togan, Torsten Günther, Füsun Özer, Eleftherios Hadjisterkotis, Mehmet Somel

**Author notes:** These authors contributed equally to this work (Torsten Günther, Füsun Özer, Eleftherios Hadjisterkotis, Mehmet Somel).

## Abstract

Once widespread in their homelands, Anatolian mouflon (*Ovis gmelini anatolica*) and Cyprian mouflon (*Ovis gmelini ophion*) were driven to near extinction during the 20th century and are currently listed as endangered populations by the IUCN. While the exact origins of these lineages remain unclear, they have been suggested to be close relatives of domestic sheep or remnants of proto-domestic sheep groups. Here, we study whole genome sequences of n=5 Anatolian mouflons and n=10 Cyprian mouflons in terms of population history and diversity, relative to eight other extant sheep lineages. We find reciprocal genetic affinity between Anatolian and Cyprian mouflons and domestic sheep, higher than all other studied wild sheep genomes, including the Iranian mouflon (*Ovis gmelini*). Despite similar recent population dynamics, Anatolian and Cyprian mouflons exhibit disparate diversity levels, which can potentially be attributed to founder effects, island isolation, introgression from domestic lineages, or different bottleneck dynamics. The lower relative mutation load found in Cyprian compared to Anatolian mouflons suggests the purging of recessive deleterious variants in the former. This agrees with estimates of a long-term small effective population size in the Cyprian mouflon. Both subspecies harbor considerable numbers of runs of homozygosity (ROH) blocks <2 Mb, which reflects the effect of small population size. Expanding our analyses to worldwide wild and feral *Ovis* genomes, we observe varying viability metrics among different lineages, and a limited consistency between viability metrics and conservation status. Factors such as recent inbreeding, introgression, and unique population dynamics may contribute to the observed disparities.

## Introduction

The Anthropocene has been an era of major shifts in species distributions worldwide. With the ongoing increase of human activity, excessive land use and resource exploitation are leading to an unprecedented acceleration in extinction rates. Over 40,000 species are currently considered at extinction risk (IUCN 2022), with the 6th mass extinction thought to be underway [1]. The International Union for Conservation of Nature (IUCN) assesses current extinction risk status of species using metrics of geographic range and recent population size change estimates. Although the genetic diversity of populations and individuals is also expected to influence extinction risk, genetic information is not included in IUCN assessments, such that populations with similar IUCN metrics can differ significantly in their genetic diversity and structure [2,3]. This has led to calls for closer integration of genetics with conservation assessment [3,4].

A major phenomenon conservation genetics centers on is the process of genome erosion with population decline. Small and fragmented populations become more prone to the detrimental effects of genetic drift and inbreeding, leading to A and F type extinction vortices, which are characterized by an increase in the species’ deleterious mutation load and a decrease in the adaptive potential, respectively [5,6]. These extinction-related dynamics can be reflected as a reduction in heterozygosity, elevated P_n_/P_s_ ratios, and long runs of homozygosity (ROH) segments. However, factors such as different demographic histories or generation intervals can produce differing levels and patterns of genome erosion signals among declining taxa [7]. For instance, depending on the nature of population decline, such as sudden bottlenecks, repeated founder effects or long-term small population size, one may or may not observe a deflated Pn/Ps ratio due to the purging of deleterious genetic load [8]. Therefore, assessing population viability can be challenging also using genetic parameters.

The Cyprian mouflon (*Ovis gmelini ophion*) and Anatolian mouflon (*Ovis gmelini anatolica*) are both endemic taxa and both have undergone bottlenecks over the last few decades, rendering them vulnerable to the detrimental effects of genetic drift and inbreeding [9–14]. Both mouflon taxa are genetically and phenotypically closely related and are identified as two subspecies of *Ovis gmelini* [15]. The latter is currently represented by three to five subspecies ranging through Armenia, Iraq, Iran, Turkey and Cyprus [16]. Unlike the “Near Threatened” status assigned to the whole species, both *O. g. ophion* and *O. g. anatolica* are considered to be “Endangered” due to their small population sizes and continuing decline [12]. The Cyprian mouflon is also listed in the Convention on International Trade in Endangered Species of Wild Fauna and Flora (CITES) appendix [12].

The history and population origins of the Anatolian and Cyprian mouflons are not well understood. Zooarchaeological evidence of sheep in Southwest Asia prior to the Holocene (∼10,500 BP) points to Central/Eastern Anatolia as a putative region of domestication [17,18]. The Anatolian mouflon may be a sister clade to domestic sheep or a feral relic of these ancient domesticates [19,20]. Meanwhile, the earliest zooarcheological record of sheep on the island of Cyprus dates back to ∼10,000 BP [18] (Supp. Note 1). These sheep possibly originate from Anatolia or the Levant; they are thought to have been brought at the early stages of sheep domestication and later feralised [21–24].

Once widespread in their homelands, due to excessive hunting, poaching, habitat loss, niche overlaps with domestic sheep/goat and predation by stray dogs [11] (Supp. Note 1), both subspecies were driven to near extinction in the 20th century. The populations experienced intense bottlenecks, with the number of individuals reduced to ∼40 in the 1930s (CYM) and ∼50 in the 1960s (ANM) [10,12,25–29]. The current population sizes have rebounded and are estimated to be 2500-3000 (CYM) and ∼900 (ANM) [12,30] . The Anatolian mouflon is found at 8 small sites, 7 of which are reintroduction sites, while the Cyprian mouflon is confined to only one reserve [10,26]. Both subspecies are legally protected, but seasonally regulated hunting has been permitted in Turkey in the recent past.

Here, we study the population history and diversity in the endemic Anatolian and Cyprian mouflon populations using published and newly generated genomic data. Joining this data with data from eight other extant sheep lineages, we investigate divergence and gene flow, as well as various metrics of diversity, population size estimates, and mutation load. We specifically ask whether the genetic diversity landscapes of Anatolian and Cyprian sheep are shaped by differences in mainland *vs*. island dynamics (e.g. isolation as well as the absence of natural predators and competition), or whether recent severe bottlenecks and genome erosion may have created similar diversity landscapes. In addition to this question, we further compare these metrics and IUCN status among the studied sheep lineages.

## Results

We generated whole genome data for n=8 Cyprian mouflon samples with a median coverage of 2.6x (ranging from 1.8x to 17.4x). In combination with publicly available n=2 Cyprian mouflon (CYM) and n=5 Anatolian mouflon (ANM), n=6 Asiatic mouflon (ASM), n=3 European mouflon (EUM), n=6 Argali sheep (ARG), n=6 Bighorn sheep (BGH), n=6 Snow sheep (SNW), n=5 Thinhorn sheep (THN), n=4 Urial sheep (URI) and n=6 domestic sheep (DOM) samples, we analyzed n=57 sheep genomes in total (Figure 1A, Table 1) [31–34]. The 6 domestic breeds were from Asia, the Middle East, Europe, and Africa, chosen to reflect the global diversity of this group. Sequence reads were mapped to the domestic sheep reference genome Oar_v4.0, genome-wide coverages ranging between 0.4x-17.4x (median = 7.5x) (Table 1).

**Figure 1.**
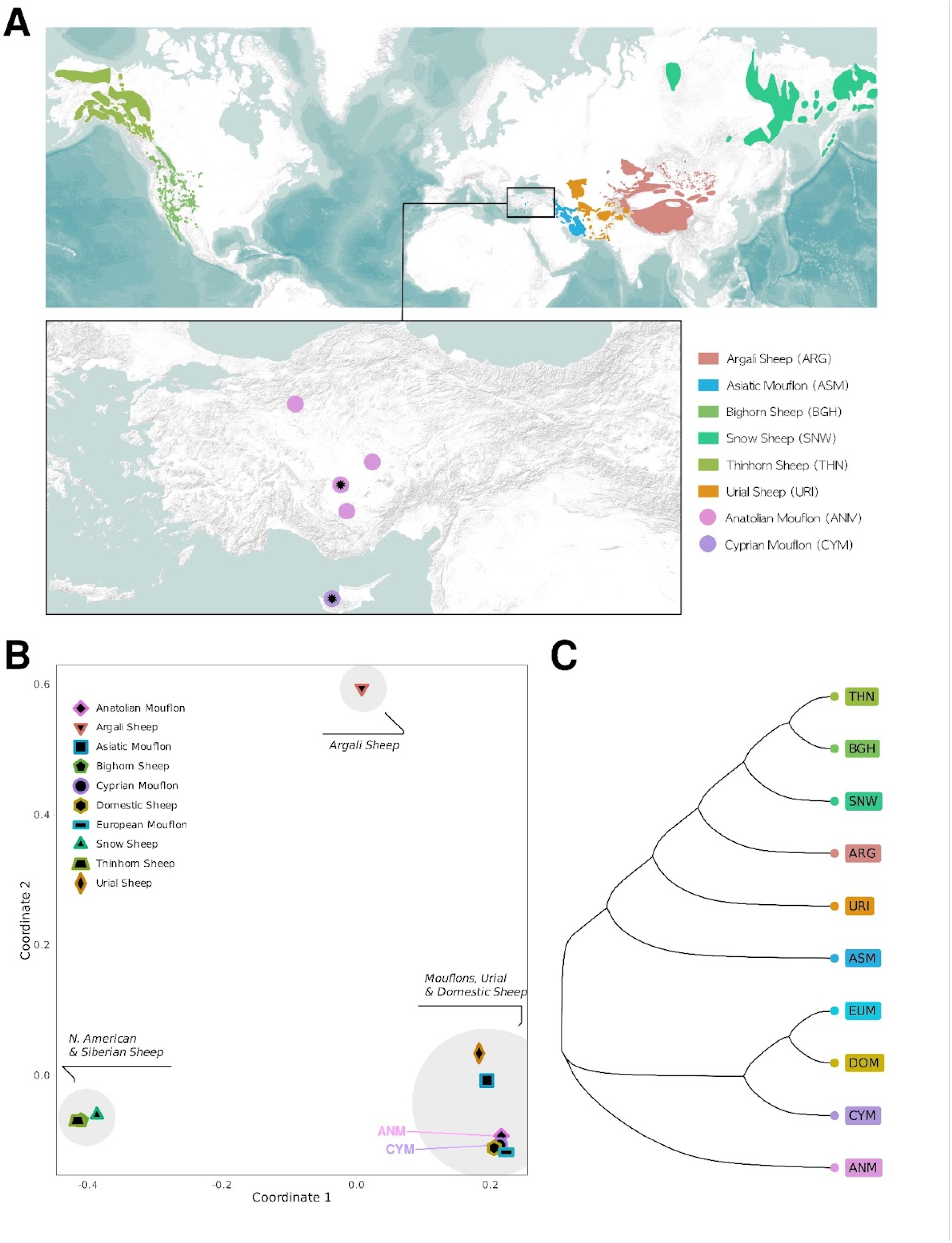
Geographic distribution of the studied genomes and the phylogenetic relationships between these sheep lineages. (A) Geographic distribution of the sheep species considered as wild by IUCN (data from IUCN). The distribution ranges of the newly generated Cyprian and Anatolian mouflons are shown as points. The points with the black symbols in the middle denote the sampling locations. (B) Multidimensional scaling (MDS) analysis of the studied sheep lineages, using 1- outgroup *f_3_* statistics as distance proxy. (C) Neighbour joining (NJ) tree of the studied sheep lineages, using (1 - outgroup *f_3_*) as distance proxies. This tree was observed in 80% of the 500 bootstraps.

**Table 1.** General information on the newly generated and published genomes.

| Sample ID | Lineage | Taxonomy | Region | Coverage | Sex | Source | Project IDs |
| --- | --- | --- | --- | --- | --- | --- | --- |
| cym002 | Cyprian Mouflon | *Ovis gmelini ophion* | Cyprus | 2.61590 | XY | This study | PRJEB69690 |
| cym003 | Cyprian Mouflon | *Ovis gmelini ophion* | Cyprus | 3.20303 | XY | This study | PRJEB69690 |
| cym004 | Cyprian Mouflon | *Ovis gmelini ophion* | Cyprus | 4.14295 | XY | This study | PRJEB69690 |
| cym006 | Cyprian Mouflon | *Ovis gmelini ophion* | Cyprus | 2.62916 | XX | This study | PRJEB69690 |
| cym007 | Cyprian Mouflon | *Ovis gmelini ophion* | Cyprus | 2.55918 | XY | This study | PRJEB69690 |
| cym008 | Cyprian Mouflon | *Ovis gmelini ophion* | Cyprus | 17.43510 | XY | This study | PRJEB69690 |
| cym009 | Cyprian Mouflon | *Ovis gmelini ophion* | Cyprus | 2.54698 | XX | This study | PRJEB69690 |
| cym011 | Cyprian Mouflon | *Ovis gmelini ophion* | Cyprus | 2.91469 | XX | This study | PRJEB69690 |
| cym012 | Cyprian Mouflon | *Ovis gmelini ophion* | Cyprus | 2.70197 | XX | Kaptan et. al, 2023 in prep. | PRJEB69690 |
| cym013 | Cyprian Mouflon | *Ovis gmelini ophion* | Cyprus | 1.80084 | XY | Kaptan et. al, 2023 in prep. | PRJEB69690 |
| OGA009 | Anatolian Mouflon | *Ovis gmelini anatolica* | Turkey | 0.69070 | XX | Kaptan et. al, 2023 in prep. | PRJEB69690 |
| OGA014 | Anatolian Mouflon | *Ovis gmelini anatolica* | Turkey | 1.02032 | XX | Kaptan et. al, 2023 in prep. | PRJEB69690 |
| oga018 | Anatolian Mouflon | *Ovis gmelini anatolica* | Turkey | 15.59840 | XX | Kaptan et. al, 2023 in prep. | PRJEB69690 |
| OGA021 | Anatolian Mouflon | *Ovis gmelini anatolica* | Turkey | 0.55503 | XX | Kaptan et. al, 2023 in prep. | PRJEB69690 |
| OGA022 | Anatolian Mouflon | *Ovis gmelini anatolica* | Turkey | 0.40399 | XX | Kaptan et. al, 2023 in prep. | PRJEB69690 |
| YZ.11 | Asiatic Mouflon | *Ovis gmelini* | Iran | 14.82340 | XX | Li et al., 2020 | PRJNA624020 |
| YZ.12 | Asiatic Mouflon | *Ovis gmelini* | Iran | 11.91190 | XX | Li et al., 2020 | PRJNA624020 |
| TH.1 | Asiatic Mouflon | *Ovis gmelini* | Iran | 14.61810 | XY | Li et al., 2020 | PRJNA624020 |
| KR.6 | Asiatic Mouflon | *Ovis gmelini* | Iran | 13.48490 | XY | Li et al., 2020 | PRJNA624020 |
| 266 | Asiatic Mouflon | *Ovis gmelini* | Iran | 12.36970 | XY | Li et al., 2020 | PRJNA624020 |
| SH-7 | Asiatic Mouflon | *Ovis gmelini* | Iran | 11.10100 | XY | Li et al., 2020 | PRJNA624020 |
| MUF1 | Europe Mouflon | *Ovis aries musimon* | Finland | 7.98631 | XX | Chen et al., 2021 | PRJNA764308 |
| MUF2-1 | Europe Mouflon | *Ovis aries musimon* | Finland | 8.37399 | XX | Chen et al., 2021 | PRJNA764308 |
| MUF3-1 | Europe Mouflon | *Ovis aries musimon* | Finland | 7.31883 | XX | Chen et al., 2021 | PRJNA764308 |
| ARG20 | Argali Sheep | *Ovis ammon* | China | 7.39554 | XY | Deng et al., 2020 | PRJNA645671 |
| ARG3-1 | Argali Sheep | *Ovis ammon* | China | 7.92511 | XY | Deng et al., 2020 | PRJNA645671 |
| ARG23 | Argali Sheep | *Ovis ammon* | China | 7.54402 | XY | Deng et al., 2020 | PRJNA645671 |
| ARG8-2 | Argali Sheep | *Ovis ammon* | China | 6.73135 | XY | Deng et al., 2020 | PRJNA645671 |
| ARG9-3 | Argali Sheep | *Ovis ammon* | China | 7.89590 | XY | Deng et al., 2020 | PRJNA645671 |
| KS1 | Argali Sheep | *Ovis ammon polii* | China | 15.72970 | XY | Yang et al., 2017 | PRJNA391748 |
| bighorn1 | Bighorn Sheep | *Ovis canadensis* | Canada | 7.31009 | XY | Chen et al., 2021 | PRJNA764308 |
| bighorn2 | Bighorn Sheep | *Ovis canadensis* | Canada | 8.15796 | XX | Chen et al., 2021 | PRJNA764308 |
| bighorn3 | Bighorn Sheep | *Ovis canadensis* | Canada | 8.13203 | XY | Chen et al., 2021 | PRJNA764308 |
| bighorn4 | Bighorn Sheep | *Ovis canadensis* | Canada | 8.13245 | XX | Chen et al., 2021 | PRJNA764308 |
| bighorn5 | Bighorn Sheep | *Ovis canadensis* | Canada | 7.87818 | XY | Chen et al., 2021 | PRJNA764308 |
| Ovican1 (BS48) | Bighorn Sheep | *Ovis canadensis* | - | 8.30279 | XY | Broad Institute | PRJNA399410 |
| snow7 | Snow Sheep | *Ovis nivicola* | Siberia | 6.83445 | XX | Chen et al., 2021 | PRJNA764308 |
| snw1 | Snow Sheep | *Ovis nivicola* | Siberia | 6.96879 | XX | Chen et al., 2021 | PRJNA764308 |
| snow13 | Snow Sheep | *Ovis nivicola* | Siberia | 6.74061 | XY | Chen et al., 2021 | PRJNA764308 |
| snow5 | Snow Sheep | *Ovis nivicola* | Siberia | 7.74575 | XY | Chen et al., 2021 | PRJNA764308 |
| snow6 | Snow Sheep | *Ovis nivicola* | Siberia | 7.21846 | XX | Chen et al., 2021 | PRJNA764308 |
| snow8 | Snow Sheep | *Ovis nivicola* | Siberia | 8.08442 | XY | Chen et al., 2021 | PRJNA764308 |
| Thinhorn1 | Thinhorn Sheep | *Ovis dalli* | Canada | 8.52089 | XY | Chen et al., 2021 | PRJNA764308 |
| Thinhorn3 | Thinhorn Sheep | *Ovis dalli* | Canada | 9.57163 | XY | Chen et al., 2021 | PRJNA764308 |
| Thinhorn4 | Thinhorn Sheep | *Ovis dalli* | Canada | 8.42533 | XY | Chen et al., 2021 | PRJNA764308 |
| Thinhorn5 | Thinhorn Sheep | *Ovis dalli* | Canada | 8.09971 | XY | Chen et al., 2021 | PRJNA764308 |
| Thinhorn9 | Thinhorn Sheep | *Ovis dalli* | Canada | 7.51948 | XY | Chen et al., 2021 | PRJNA764308 |
| BJ3-1 | Urial Sheep | *Ovis vignei* | Iran | 6.86997 | XX | Chen et al., 2021 | PRJNA764308 |
| BJ4-1 | Urial Sheep | *Ovis vignei* | Iran | 7.52407 | XX | Chen et al., 2021 | PRJNA764308 |
| BJR5 | Urial Sheep | *Ovis vignei* | Iran | 6.90093 | XX | Chen et al., 2021 | PRJNA764308 |
| BJ1-2 | Urial Sheep | *Ovis vignei* | Iran | 6.82588 | XX | Chen et al., 2021 | PRJNA764308 |
| AL118 | Altay sheep | *Ovis aries* | China | 11.72370 | XX | Li et al., 2020 | PRJNA624020 |
| FINN305 | Finnsheep | *Ovis aries* | Finland | 12.71830 | XX | Li et al., 2020 | PRJNA624020 |
| Sc12 | Mbororo sheep | *Ovis aries* | Cameroon | 16.66190 | XY | Li et al., 2020 | PRJNA624020 |
| DLS249 | Duolang | *Ovis aries* | China | 11.28540 | XX | Li et al., 2020 | PRJNA624020 |
| CC50 | Cine-Capari | *Ovis aries* | Turkey | 6.43194 | XY | Kijas et al., 2012 | PRJNA160933 |
| AWA-33 | Awassi | *Ovis aries* | Iraq | 7.19223 | XY | Deng et al., 2020 | PRJNA645671 |
| BAT_IOSW | Goat | *Capra hircus* | Morocco | 5.01279 | XY | NextGen project | PRJEB3134 |

Genetic sex was determined with the *R_x_* metric (Table S2). All 5 ANM and 4 of the 10 CYM individuals were female. We found one pair of genetically identical genomes (possible twins or sample duplicates) (Table S3). One individual from this pair was excluded from the analyses to ensure the independence of the sample. We also found one pair of possible 2nd-degree and eight pairs of possible 3rd-degree relative pairs, amongst which we excluded two individuals (Methods).

In order to study variation, we created a SNP dataset involving all the sheep populations. We called SNPs using one representative genome from each lineage with similar data quality. Specifically, we chose one individual for each of the 10 populations and downsampled these to similar coverages, 7.5-8.5x (Table 1, S1), and then performed *de novo* SNP calling on each and combined the data. In the end, we obtained a sheep variation dataset comprising ∼14 million autosomal SNPs (Methods). The dataset was used for the calculation of f-statistics, and identification of ROH segments and biologically related individuals. We note that this approach limits ascertainment bias compared to calling SNPs from the full dataset, but does not fully eliminate such bias as the lineages are not equally distant to each other. Nevertheless, we also validated our main findings possibly prone to ascertainment bias using a ∼116k subset of the ∼14M SNPs that were identified as heterozygous in a goat individual (we lacked an outgroup phylogenetically close enough for large-scale SNP ascertainment) (Methods).

### Demography

To summarize the differentiation between the populations, we calculated genetic distances using 1 - outgroup-*f_3_*statistics (Table S3). Amongst the studied sheep populations, ANM and CYM were most distant to populations from North America and Siberia, and showed the highest affinity to European mouflon and domestic sheep. Employing Multidimensional Scaling (MDS) with 1 - outgroup-*f_3_* statistics as distance proxies, we observed 3 separate clusters (Figure 1B, S1). North American and Siberian wild sheep comprised one cluster and the Argali sheep another, positioned separately from all other sheep. In the third cluster, Cyprian and Anatolian mouflons grouped with the other mouflons, as well as Urial and domestic sheep. We also utilized 1-outgroup *f_3_* values to construct a Neighbour Joining (NJ) tree with 500 bootstraps. In the majority tree (80% bootstrap support), whilst the Anatolian mouflon diverged as a separate lineage, Cyprian mouflon formed a clade with the European mouflon and domestic sheep (Figure 1C, S2). We further validated the clustering in the MDS and the structure of the NJ tree with the dataset of goat heterozygous SNPs (Figure S3). The above pattern is also observed in the comparison of *f_3_* and *f_4_* values with wild *vs*. domestic sheep, demonstrating that domestic sheep and European mouflons (which are considered feral populations of past domestic livestock) show higher affinity to the Cyprian mouflon than ANM, and reciprocally, the Cyprian mouflon is closer to domestic sheep and European mouflons than to ANM (Figure S4-5, Table S4-5). Meanwhile, both ANM and CYM appeared to share a similar amount of drift with other wild sheep lineages.

We also analyzed the mitochondrial DNA haplogroup distribution by using median-joining (MJ) network. This revealed 3 distinct branches (Figure S6). BGH, SNW and THN were clustered on one branch, meanwhile, ARG, URI and 3 ASM individuals clustered together in another branch. The last branch was composed of DOM, CYM, ANM, EUM and the remaining 3 ASM individuals. Within this third clade, domestic sheep were clustered into five different haplogroups (hpg) named A-E [35]. EUM samples were clustered with hpgB (the most widely observed haplogroup in domestic sheep of Europe). Meanwhile, ANM clustered in two different sub-branches, n=3 individuals (OGA009, oga018, OGA022) were on hpgA and n=2 individuals (OGA014, OGA021) were on a distinct branch near hpgC and hpgE, named hpgX [19]. Finally, all CYM samples were clustered near hpgX. The patterns differ from the autosomal clustering in the sense that CYM clusters with ANM rather than DOM and EUM, which suggests different population histories on the maternal line.

The ANM and CYM populations are thought to have undergone severe population declines in the recent past, which should leave detectable signatures in their genomic diversity data. Employing the Pairwise Sequential Markovian Coalescent (PSMC) approach to the highest coverage individuals (cym008 & oga018), we estimated the change in historical effective population sizes (N_e_). The demographic trajectory of the CYM shows an extended period of reduction in N_e_ (Figure 2). There seems to have been periodic declines of a similar extent, followed by a stabilization near 10 kya. Meanwhile, ANM shows a period of population growth starting from 50 kya, followed by a sudden decline near 10 kya. Considering the sensitivity of the PSMC approach to genome-wide coverage [36], we also ran a trial with samples downsampled to similar coverages (7.5-8.5x). Although the trajectories inferred from the original and downsampled data were partly different, the main qualitative patterns were similar: we still observed a period of population growth in ANM and a substantially long period of decline in CYM (Figure S7). In both results, compared with the most recent N_e_ values estimated from individuals of DOM and other wild sheep populations, CYM shows the lowest estimate.

**Figure 2.**
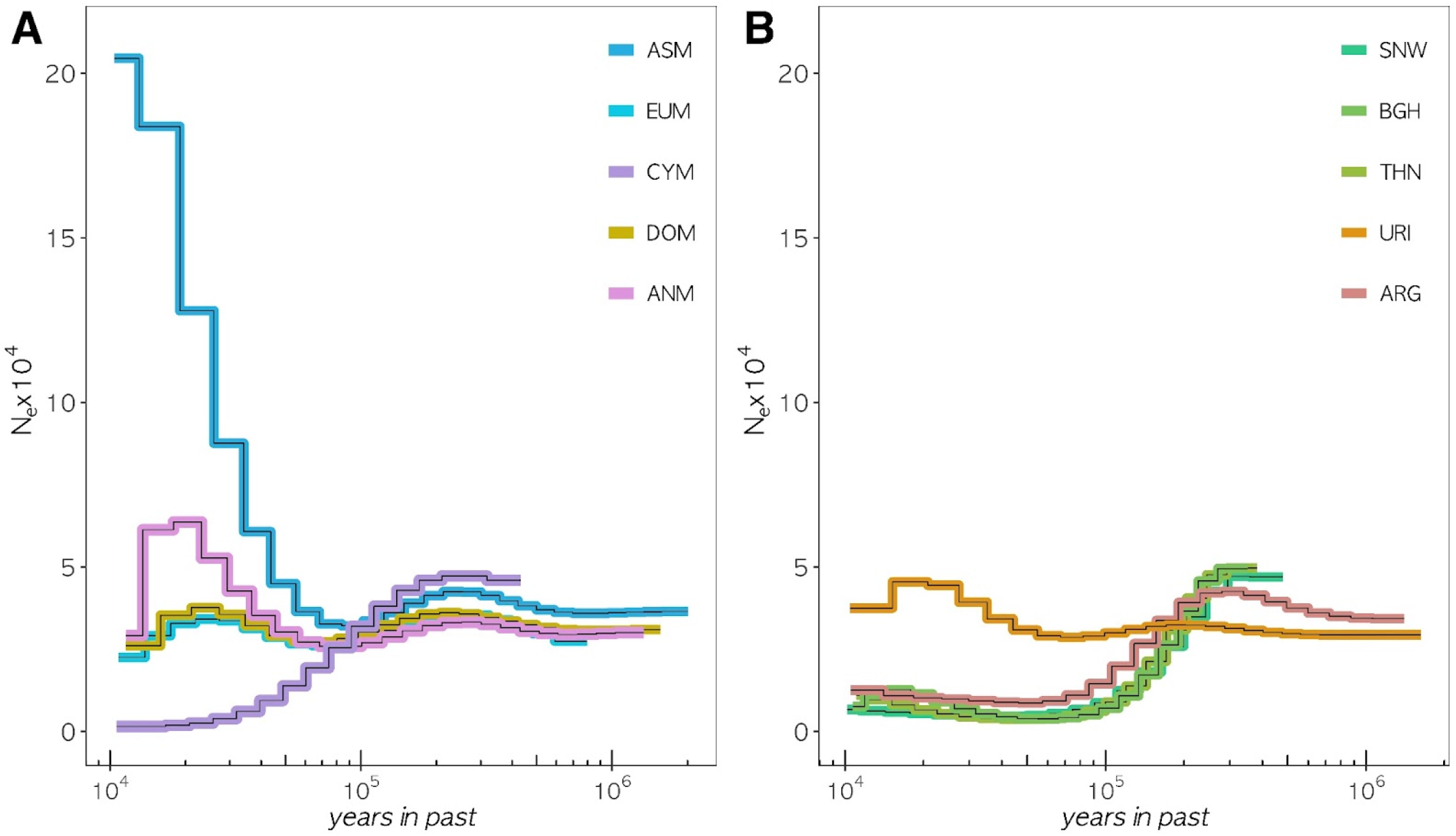
Population history of the studied sheep lineages. PSMC analysis of high coverage individuals from each lineage, (A) for mouflons and domestic sheep, and (B) for N. American, Siberian and Asian wild sheep. The x-axis shows time in a log scale, and the y-axis shows the estimated effective population size. We assumed a generation time of 3 years and a mutation rate of 1.5×10^-8^.

### Diversity

Next, we studied within-population population diversity patterns in CYM and ANM and compared these with the sheep lineages. For this, we used both genome-wide heterozygosity (π) and inter-individual diversity estimates using pairwise 1- outgroup-*f_3_* statistics (Figure 3). For π, we used the highest coverage individuals (cym008 & oga018) for CYM and ANM, and ran the analyses on genomes downsampled to similar coverages (6.5 - 7.5x). The North American/Siberian group had the lowest π followed by the European mouflon (Figure 3A, Table S6). The Cyprian and Anatolian mouflon recorded moderate and high estimates, respectively. The highest value was observed for the Urial sheep, more than 4 times higher than the North American/Siberian sheep.

**Figure 3.**
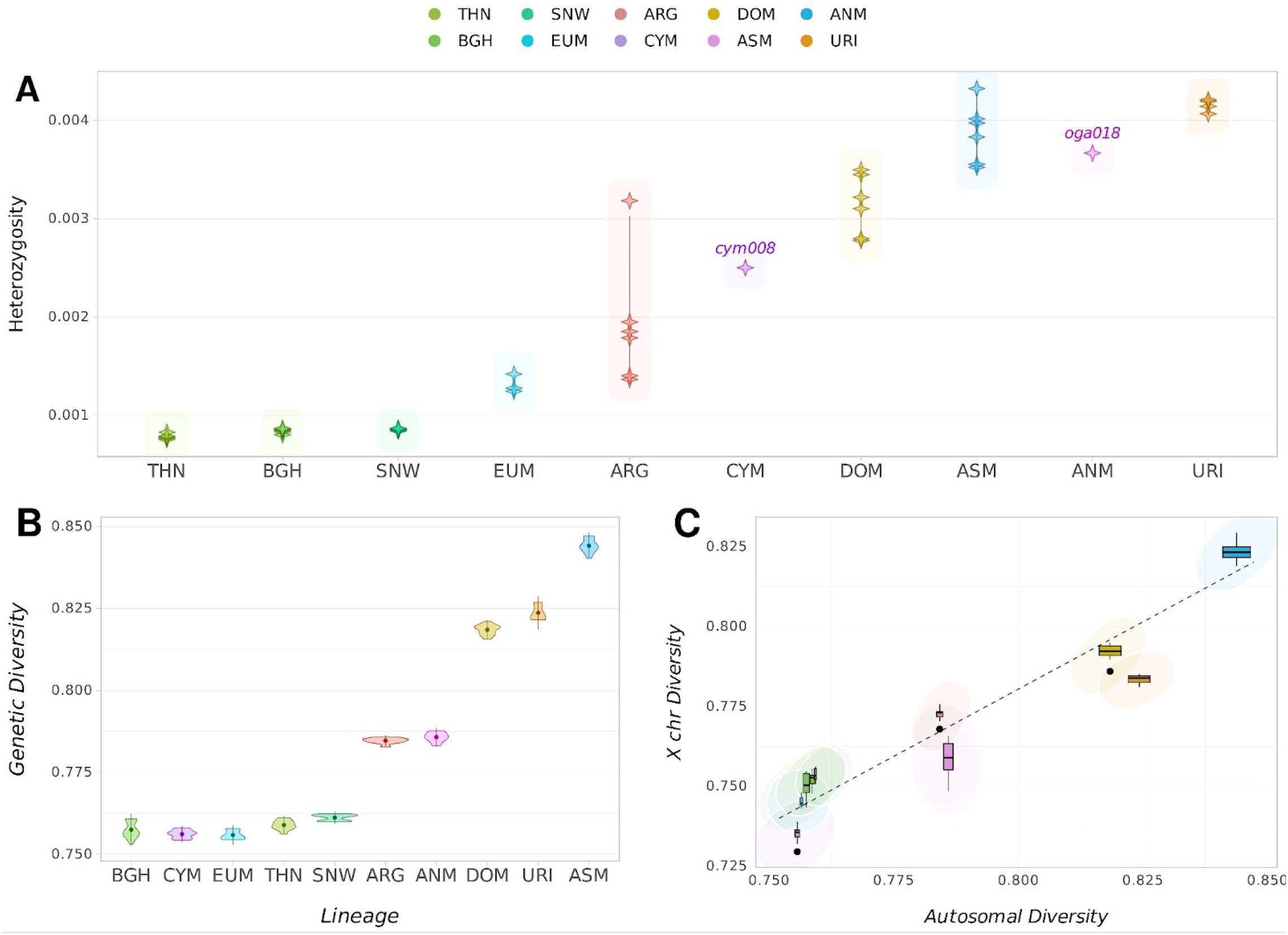
Heterozygosity and diversity estimates of the studied sheep lineages. (A) Genome-wide heterozygosity values estimated using genotype likelihoods. Only the high coverage individuals cym008 and oga018 from the ANM and CYM populations were included. (B) Within-population diversity values estimated using pairwise 1 - outgroup *f_3_* statistics per lineage. (C) Comparison of autosomal *vs*. X chromosome diversities, each estimated using pairwise 1 - outgroup *f_3_* statistics. The regression line was generated with the loess algorithm in the R *stats* package.

In order to include the low-coverage CYM and ANM individuals in the analyses, we then estimated inter-individual diversities using pairwise 1- outgroup-*f_3_* statistics between individuals within each population as a proxy for population-wide heterozygosity (Figure 3B). The highest inter-individual diversity was observed for the Asiatic mouflon, which is thought to have experienced gene flow from other wild and domestic sheep populations [31,37]. Cyprian mouflon showed one of the lowest diversity estimates along with the European mouflon and the North American/Siberian group. The Anatolian mouflon was found to have a relatively moderate level of inter-individual diversity. We observed qualitatively similar results using the dataset of goat heterozygous SNPs (Figure S3).

We further studied mitochondrial and X chromosomal diversities across these genomes (Figure S8-10). These showed similar patterns to autosomal diversity estimates, with CYM showing the lowest and ANM moderate values compared to other sheep.

We then compared X chromosome diversities with autosomal diversities across the ten lineages (Figure 3C, S9). The autosome / X ratios ranged between 1.01 and 1.05, lower than the expected proportion of 1.33 assuming equal N_e_’s for both sexes (Figure S9). These values suggest smaller male N_e_, consistent with the harem-like mating structure in sheep However, the lineages also varied among themselves: we found that the autosomal / X diversity ratios were closest to 1 in the N. America / Siberian group, followed by the European mouflon and Argali sheep. This may be compatible with a relatively higher female contribution to the genetic variation in these populations, i.e. a smaller relative male N_e_. In contrast, other populations including CYM and ANM showed relatively lower X chromosomal diversities, implying that male *vs*. female N_e_ differences may be more modest in these groups.

### Inbreeding

In order to measure inbreeding levels across these 10 sheep lineages, we studied Runs of Homozygosity (ROH) segments >500kb found in individuals’ genomes. CYM and ANM genomes were amongst those with a high ROH load, in terms of both the total number and size of the called segments (Figure 4A, Table S7). Between the two, CYM had a higher load, in line with its relatively depleted diversity. We next studied the relative frequencies of four different ROH size classes, (0.5 - 1 Mb), (1 - 2 Mb), (2 - 3 Mb) and (3 - 5 Mb), which we used to estimate the time to the most recent common ancestor corresponding to each class in relation to recombination rate and generation time [38,39]. We had four time frames spanning from 200 years ago to 20 years ago, assuming a generation time of 3 years (Figure 4B). ANM and CYM had a mean ROH length of 0.63 Mb and 0.82 Mb, suggesting a common ancestor 53 and 41 generations ago, respectively (121 and 157 years ago). Neither carried segments longer than 3 Mb and had relatively low proportions of segments of 2nd and 3rd class; this result suggests bottlenecks and small historical population size as sources of ROH rather than recent inbreeding. We also calculated the proportion of the genomes harboring ROH segments, referred to as F_ROH_ (Figure 4C). Fourteen per cent of the ANM genome contained ROH segments, while CYM had a higher F_ROH_ of 20%, preceded by the European Mouflon, which had the highest estimate of 42%.

**Figure 4.**
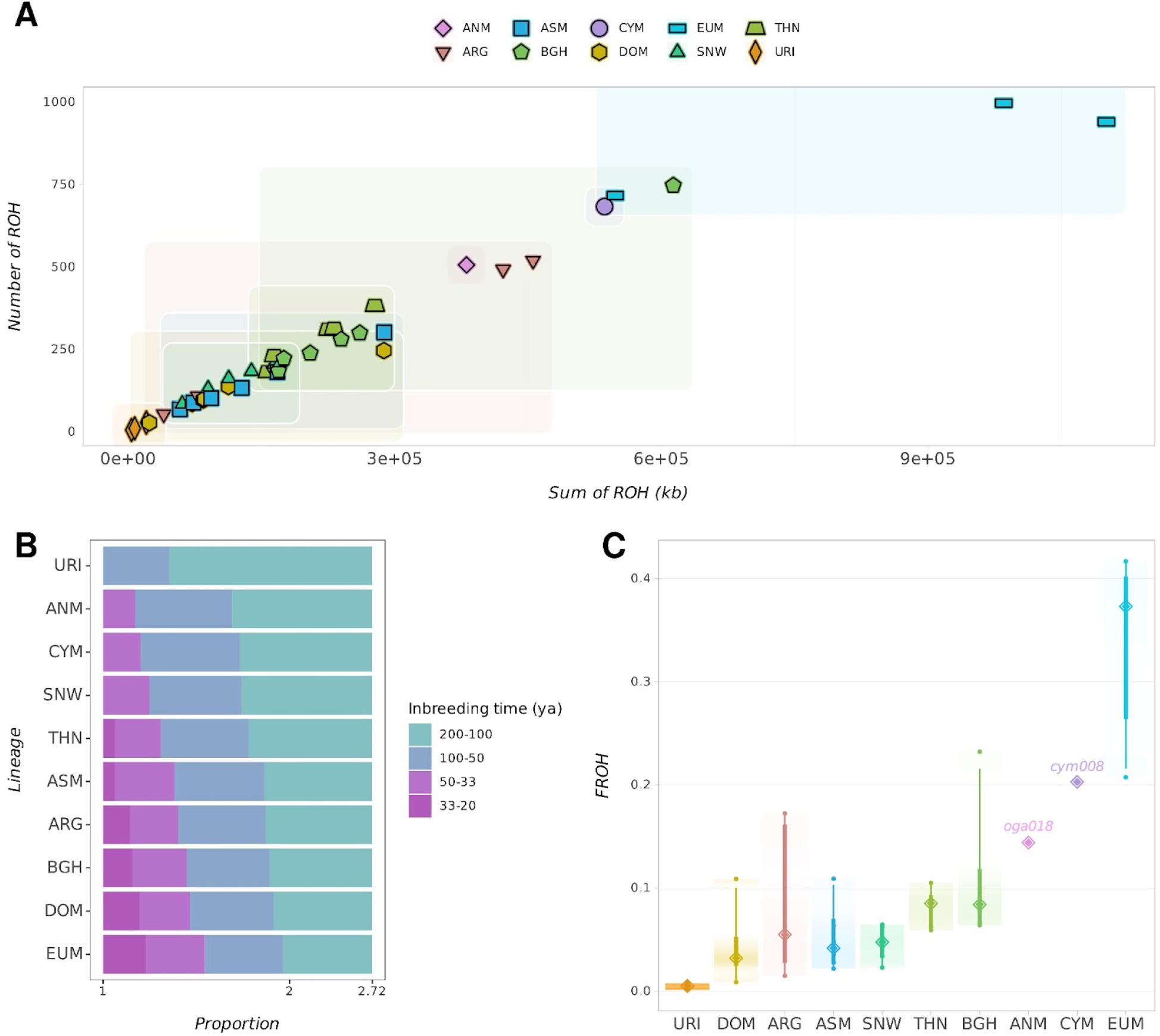
Runs of Homozygosity (ROH) analyses. (A) Number of ROH segments longer than 500 kb plotted against the total length of the segments found in each individual. The ANM and CYM populations are only represented by the high-coverage individuals oga018 and cym008, respectively. (B) Size distribution of ROH segments divided into four classes (0.5-1 Mb, 1-2 Mb, 2-3 Mb, 3-5 Mb). Inbreeding times corresponding to each size class was estimated assuming a generation time of 3 years and a recombination rate of 1.5 cM/Mb [38,39]. The x-axis is given in log scale. (C) Proportion of ROH segments longer than 500kb in each individual’s genome.

### Mutation load

We further tested possible elevations in mutation load due to historic bottlenecks and small population sizes in CYM, ANM, and the other 8 sheep lineages. For this, we assessed the substitutions in evolutionary conserved genomic regions, utilizing Genomic Evolutionary Rate Profiling (GERP) scores [40]. We chose stretches of sites with GERP scores >4 as highly conserved regions. The relative mutation load for each sample was estimated by calculating the normalized GERP scores for the derived alleles observed in the conserved regions [41].

The load estimates showed substantial variation among individuals from the same taxon. Nevertheless, we did observe systematic patterns, with the lowest average load estimates amongst the Asiatic mouflon, and the wild sheep from N. America and Siberia harboring the highest values (Figure 5). Interestingly, CYM and ANM genomes had low-to-moderate relative mutation load estimates, with a slightly lower estimate for CYM. The discrepancies between degrees of diversity and mutation load may stem from differences in the duration and timing of the population bottlenecks (see Discussion).

**Figure 5.**
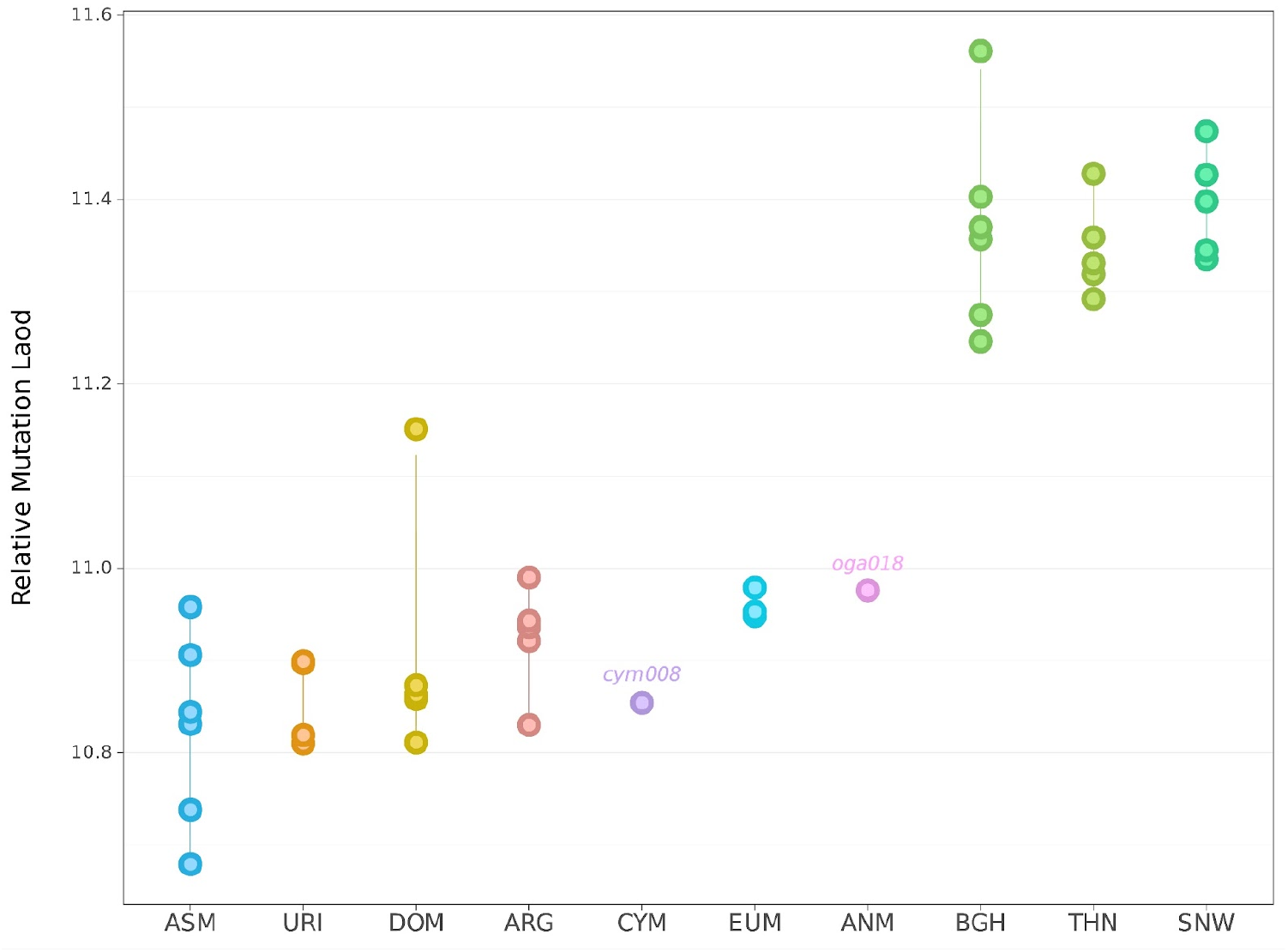
Mutation load estimates using GERP scores. Relative mutation loads were calculated as average GERP scores weighted by the number of derived variants, only including variants found in conserved regions (GERP > 4) [41]. We used goat alleles to infer the derived state. Only the high-coverage individuals cym008 and oga018 from the ANM and CYM populations were included.

### Co-evaluation of viability metrics and conservation status

Finally, we assessed the conservation status of each population in relation to their studied viability metrics. DOM and EUM were excluded since these two are not evaluated by the IUCN. First, we compared the individual heterozygosity estimates with the F_ROH_ values (Figure 6A). We did not find a significant correlation between the two metrics (p=0.113). On the other hand, the relationship between the relative mutation load and heterozygosity was stronger (ρ=-0.869, p=1.1e-10). Second, we compared the relationship between the IUCN status and the two viability estimates (Figure 6B). In line with previous observations [2,3], the IUCN status does not seem to be indicative of the population diversities in sheep, as “Least Concern” populations have lower heterozygosity than “Endangered”, “Near Threatened” and “Vulnerable” populations. Likewise, the “Least Concern” populations (N. America / Siberia) show the highest mutation load. We also do not observe a strong relationship between IUCN status and F_ROH_, except that the “Endangered” ANM and CYM show the highest F_ROH_ values. Various interplays between demographic events such as introgression, founder effects and isolation might explain this lack of relationship between the metrics.

**Figure 6.**
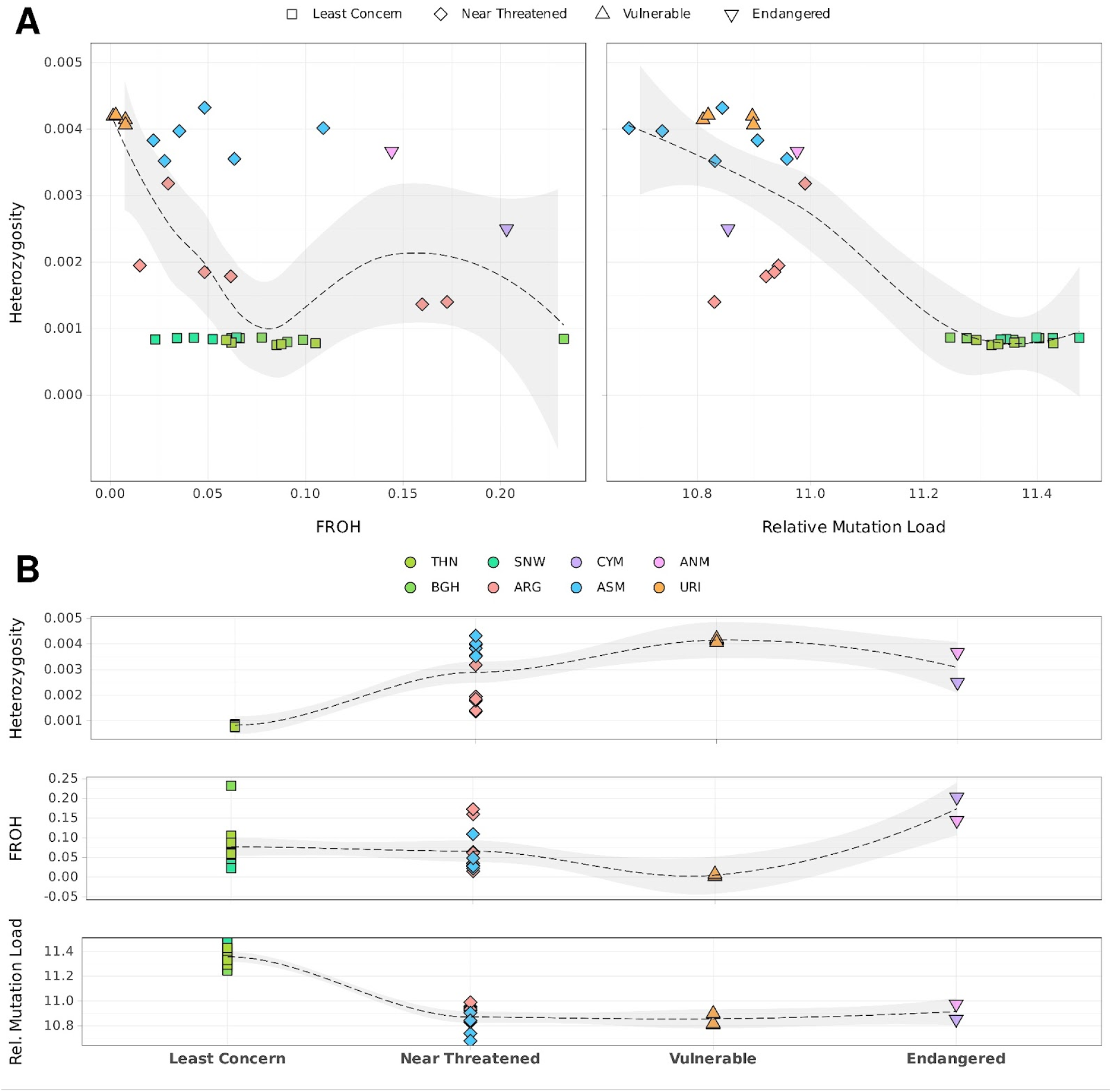
Co-evaluation of genetic viability metrics and IUCN status. (A) Correlations between viability metrics per individual compared across sheep lineages assessed by the IUCN. Regression lines were calculated using the method “loess” in the R *stats* package. (B) The IUCN status compared with the genetic viability metrics heterozygosity (π), F_ROH_, and relative mutation load.

## Discussion

### Phylogenetic relationships between Anatolian and Cyprian mouflons and domestic sheep

Previous work reported that ASM was the wild sheep lineage genetically closest to DOM, relative to other wild sheep, implying that ASM could have been the wild source of domestic sheep [42–45]. Here we found that ANM and CYM cluster with EUM and DOM, with ASM as outgroup, a pattern supported by MDS, *f_3_* and *f_4_* statistics. Further, we observed a higher affinity of DOM to CYM than to ANM. These results could be compatible with multiple scenarios: (a) The ancestors of ASM and ANM contributed equally to domestic sheep, but recent Urial introgression into ASM [31] differentiated ASM from the ANM-CYM-DOM cluster. (b) The source of domestic sheep were the ancestors of ANM and CYM but not ASM, and therefore ANM and CYM share closer ancestry with EUM and DOM than ASM. This scenario would also be compatible with the suggestion that CYM could have been a proto-domesticate lineage. The positive f_4_(Goat, DOM; ANM, CYM) result is also consistent with the notion that CYM was an early feral lineage that split from the domesticated sheep gene pool. (c) Recent DOM introgression into ANM and CYM. This is also possible given the significant positive f_4_ statistics of the form f_4_(Goat, ANM; CYM, DOM) and f_4_(Goat, CYM; ANM, DOM). We are currently unable to reject any of these scenarios. Information from ancient genomes may prove to be essential for resolving the complicated demographic history of domestic sheep [46].

### Differences in demographic history between Anatolian and Cyprian mouflons

Our data also allowed us to investigate the genomic footprints of the population size fluctuations via different viability metrics. Within-population diversity and individual-level heterozygosity estimates revealed CYM as harboring low-to-moderate, and ANM as harboring moderate-to-high diversity values relative to other sheep lineages. It is intriguing that even though both subspecies ANM and CYM experienced bottlenecks of similar extent during similar time periods [10,12–14,28], their diversity levels are visibly different. The specifics of the bottleneck, such as its duration and how much of the original population structure survived, the extent of post-bottleneck population growth, and the subsequent conservation practices may have affected these diversity level differences. In addition, both populations have a history of population fluctuations due to paratuberculosis in ANM and keratoconjunctivitis and particularly poaching in CYM [10,12–14,47–49]. Other than these more recent events, PSMC analyses suggest that shifts in the effective population sizes of ANM and CYM prior to 10 kya also follow different trajectories. Different processes might have shaped these diversity estimates, such as: a) CYM losing ancestral diversity due to founder effects during its transport to Cyprus, or b) CYM undergoing serial bottlenecks in Cyprus, being isolated on an island. Both scenarios involve long periods of high homozygosity, which may have led to purging of recessive deleterious variants in CYM [50,51] and its consequent low relative mutation load compared to ANM.

We find the highest levels of ROH load in two CYM and ANM genomes relative to all other sheep lineages, except for EUM. Moreover, higher ROH in CYM than in ANM appears in agreement with the above scenarios involving smaller ancestral population size in CYM. Here, we note that our ROH analyses involve only one genome for ANM and CYM each. In both genomes, the majority of the segments are of moderate length, suggesting that recent inbreeding is not the source of the high ROH load. Instead, the signal likely results from smaller effective population size than the particular history of the studied individuals.

### Variable diversity and mutation load patterns among wild and domestic sheep genomes

Studying the genetic viability metrics of the other sheep lineages, we found that ASM and URI had the highest heterozygosity/diversity estimates, the lowest mutation loads, and on average lower ROH segments than other sheep lineages. These patterns may be consistent with the history of introgression between ASM and URI and/or domestic introgression to ASM [37,52].

EUM had exceptionally high F_ROH_ among the studied genomes. However, this result should be taken with caution as the genomes were sampled from a population in Finland transported from Sardinia/Corsica [53]; they may have thus undergone additional founder effects in the process. Meanwhile, the six DOM genomes had genetic characteristics similar to each other, with relatively high heterozygosity, low F_ROH_ and low mutation load (except for Awassi sheep which deviated from the rest with its high F_ROH_).

The N. America / Siberia group (BGH, THN, SNW) showed systematically lower heterozygosity/diversity among all studied sheep lineages, harboured by far the highest mutation load estimates, and carried intermediate levels of ROH. These three lineages also had relatively small past population size estimates in PSMC analyses, along with CYM and ARG. It is not clear why the N. America / Siberia group and CYM show disparate patterns with respect to mutation load, despite all these lineages being attributed to long-term small effective population sizes. Here we note that since the reference genome is assembled from domestic sheep, the heterozygosity/diversity estimates might be inflated/deflated depending on the populations’ genetic proximities to domestic sheep. In wild populations with higher proximity to the domestic sheep (e.g. CYM), more diversity may be represented, while in more distant populations (e.g. BGH) the estimates can be deflated. Mutation load estimates might also be affected by the asymmetric distances to the reference genome.

### Comparison of viability metrics and conservation status

Our results reveal limited consistency between different genetic viability metrics among the studied sheep lineages. In contrast to expectation, we did not find a significant correlation between heterozygosity and the proportion of ROH segments (p=0.113). Recent inbreeding and introgression can be counted among the likely causes for the observed disparity. The time past since inbreeding might be too short to impact genome-wide diversities significantly. Admixture coupled with recent inbreeding can also elevate heterozygosity while also creating a high ROH load.

Regarding the mutation load, we find a strong negative correlation with heterozygosity (ρ =-0.852, p=6e-10). Still, deviations from the general trend can be observed, such as the Cyprian mouflon with both low heterozygosity and low mutation load. Such discrepancies may emerge depending on the nature of bottlenecks and post-bottleneck population growth, as well as the nature of mutation load, such as recessiveness and degree of deleteriousness. While long-term small population sizes may induce purging, sudden bottlenecks can lead to the accumulation of deleterious mutations. Here too, possible impacts of introgression causing discrepancies cannot be excluded, as introgression may not only reduce but also contribute to the load if the introgressing population has a load of its own [8].

We also observe differences in the X chromosome *vs.* autosomal diversities between populations, indicating lower male *vs*. female N_e_ among the 10 sheep lineages, albeit at varying levels. Lower male N_e_ can be caused by male reproductive skew. Female philopatry and male dispersal are often observed in wild sheep populations [16], which may also contribute to sex differences in reproductive success and lead to low male N_e_. Meanwhile, the extent of this spatial behavior can depend on the habitat structure shaped by natural and anthropogenic factors, which may cause the observed differences among lineages. Sex-biased introgression from domestic sheep and higher mortality of males due to natural causes or hunting pressure can also be counted among driving factors behind X chromosome *vs.* autosome diversities.

Lack of a clear relationship also prevails between each lineage’s IUCN status regarding their estimates for diversity, ROH and mutation load. Since the assessment of IUCN status is based on current/recent population viability criteria, and the genomic diversity indices mostly reflect historical population characteristics, a direct relationship may not be expected, especially if the population has undergone major demographic changes [54]. Similar to recent work on a wider range of taxa, we did not find the genetic viability metrics to be indicative of the threat status among the 8 sheep lineages evaluated by IUCN [2,3]. Strikingly, although the N. America / Siberia group shows on average lower diversity levels and a distinctly high mutation load relative to other lineages, they are listed as “Least Concern”. Species from this group have relatively high census population size estimates (IUCN 2022), but our results suggest the possible vulnerability of these populations to perturbation, such as epidemics.

Finally, we touch upon the fact that European mouflons are currently not assessed by IUCN, since they are known to be feralized descendants of domestic sheep. Conservation of feralized species has constituted an issue of discussion, such as the case of Australian dingoes. These were originally considered “vulnerable” but later discarded from the IUCN Red List as their status was revised as feral dogs, although conservation efforts for protecting some dingo populations are still being carried out [55–58]. Similarly, in the case of EUM, there have been local efforts to protect populations in Sardinia and Corsica [59–62]. These efforts are relevant given that the EUM has been in the wild for nearly 10k years (even if it may have experienced further domestic introgression after feralization). Harboring the largest amount of ROH segments among all studied sheep lineages, showing low diversity and high mutation load values, the EUM population (at least those individuals from Finland included in this study) seems to have a viability estimate lower than the officially endangered Cyprian and Anatolian mouflons. Considering the substantial domestic proximity of the CYM and ANM, we suggest a reassessment of the European mouflon’s conservation status.

## Materials and Methods

### Sample collection, DNA extraction, library preparation and sequencing

Cyprian mouflon tissue samples were collected from animals found dead at the northern part of Paphos forest, located in the Troodos Mountains, mainly near the villages Kampos and Tsakistra, under the permit of the Ministry of the Interior for scientific research. DNA extraction from tissue samples was performed using MACHEREY-NAGEL “NucleoSpin tissue” kit following the standard protocol. DNA fragmentation via sonication was carried out using Qsonica Q800R at 100% amplitude for 15’’ On and 15’’ Off at 4°C for 12 minutes. Fragmented DNA samples were quantified on Agilent Bioanalyzer 2100 to confirm an average 300 bp fragment length. If samples had an average fragment length longer than 300 bp, the sonication step was repeated. Dilutions were performed accordingly. Double indexed Illumina sequencing libraries were prepared following the Meyer & Kircher 2010 protocol [63] and sequenced on NovaSeq 6000 flowcells (NovaSeq Control Software 1.7.5/RTA v3.4.4) with a 101nt(Read1)-7nt(Index1)-7nt(Index2)-101nt(Read2) setup using the ’NovaSeqXp’ workflow in ’S4’ mode flowcell.

### DNA data processing and variant calling

Residual adapter sequences were removed using *AdapterRemoval v.2.3.1* [64]. Sequence reads were mapped to the sheep reference genome Oar_v4.0, using *BWA mem v.0.7.15* with the parameter *-M* [65]. After removing the PCR duplicates with *Picard MarkDuplicates*, reads with mapping qualities lower than 20 were discarded using *samtools v.1.9* [66]. We chose one representative individual from each sheep population (Table 1), n=10 in total, and downsampled them to similar coverages between 7.5-8.5x using *samtools v.1.9* [66] *view -s*. We carried out *de novo* SNP calling using *GATK Haplotypecaller v.4.4.0.0* [67], retaining SNPs with depths between 4x-16x and a quality of 20, also using *-- maf 0.05* and *-- hwe 0.001,* followed by the genotyping of the remaining genomes (Table 1,S1). We included only biallelic SNPs and did filtering using *bcftools v.1.18* [68] with parameters *QUAL>=20 QD>=2.0 SOR<=3.0 FS<=60.0 MQ>=40.0 MQRankSum>-12.5 ReadPosRankSum >-8.0*. After the filtering steps, the resulting autosomal dataset contained a total of 14,237,712 SNPs and the X chromosome dataset contained 427,454 SNPs. We used these sets of SNPs in the calculation of *f*-statistics, identification of autosomal ROH segments and the genetic relatedness analysis with *READ*. We also validated our main findings based on *f_3_*-statistics using a subset of our autosomal dataset. For this, we only included the sites that were heterozygous in the outgroup goat genome (Table 1), which resulted in a total of 116,613 SNPs.

### Sex determination and relatedness

We determined the genetic sex of the studied genomes utilizing the *R_x_* metric [69] thresholds optimized for sheep, using SexDetermineOar (https://github.com/mskilic/SexDetermineOar). For the relatedness estimates, we used the program *READ* which detects up to 2nd-degree relatives using pseudo-haploid genotype data [70]. The pairwise mismatch (P_0_) values were normalized using the median of the P_0_ estimates of each population. Then, normalized P_0_ values were subtracted from 1 to obtain θ (kinship coefficient) estimates. We assigned the individual pairs to their respective kinship degrees using the midpoint between the two expected θ values (θ_1_ and θ_2_) as a threshold, calculated as *(θ_1_+θ_2_)/2* [70]) (Table S3). One of the identical genomes (cym012) and the potential 2nd and 3rd-degree relatives that appeared divergent in the pairwise *f_3_*estimates (cym006 and cym007) were excluded from the diversity analyses.

### Mitochondrial DNA analyses

Mitochondrial DNA gVCF files were generated using *bcftools v.1.18* [68] with parameters *-q20 -Q20 -mV indels.* Average mitochondrial pairwise differences (π) within each population were calculated using the formula:

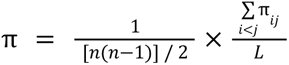

where *n* is the total number of individuals in each population, L is the total number of sites, and *π*_ij_ is the number of nucleotide differences for each pair. Only the biallelic sites which were non-missing in at least two individuals within the population were taken into account.

Mitogenome consensus sequences were generated from BAM files using *ANGSD v.0940* [71] with parameters ‘-doFasta 2’, ‘-minQ 30’, ‘-minMapQ 30’ and ‘-setMinDepth 2’. Ten representative domestic sheep mitogenomes with known haplogroups (two for each haplogroup) with NCBI-Genbank Accession No: HM236174-83 [35] were also added to the dataset. Consensus sequences were aligned with *MAFFT v.7.490* [72] and a median-joining (MJ) network [73] was constructed with *PopART* [74].

### Outgroup-*f*_3_ and *f*_4_ statistics

We calculated outgroup-*f_3_* and *f_4_* statistics using *Admixtools v.2.0.0* [75] with default parameters and *maxmiss=1* (includes all SNPs). We used goat as the outgroup (Table 1). *f_3_* statistics of the form *f_3_*(Goat,ind1,ind2) was performed for the estimation of within-population (inter-individual) genetic diversities. *f_3_*(Goat,pop1,pop2) was calculated for estimating population differentiation, pop1 and pop2 corresponding to different wild sheep populations. When comparing populations we preferred outgroup-*f_3_*instead of F_ST_ because the latter is sensitive to population size fluctuations and consequent variation in within-population diversity, while the former is not [76]; *f_3_* can therefore capture population divergence and admixture more effectively than F_ST_ [77].

To summarize the genetic relationships among populations, we used pairwise 1 - outgroup-*f_3_* values to construct a distance matrix, which we used to perform multidimensional scaling analysis (MDS) and also construct a neighbour-joining (NJ) tree. MDS was run using the function *cmdscale* implemented in R package *stats*. For the NJ tree, the function *nj* in the R package *ape v.5.7-1* [78] was utilized. We performed n=500 bootstraps by first dividing the genotype data into chunks of 30,000 SNPs, randomly sampling chunks with replacement and calculating outgroup *f_3_* statistics per iteration.

### Demographic history reconstruction

We used the Pairwise Sequential Markovian Coalescent (PSMC) model [79] to infer the changes in historical effective population sizes. PSMC was run for the highest coverage CYM and ANM individuals with parameters –N25 –t15 –r5 –p ‘4 + 25*2 + 4 + 6, using a generation time of 3 years, and a mutation rate of 1.51 × 10e-8 for scaling [80–82]. We also ran PSMC with the same genomes downsampled to similar coverages (7.5-8.5X).

### Runs of Homozygosity (ROH) and heterozygosity estimates

We used *ANGSD v.0.940* [71] for the estimation of genome-wide heterozygosities, with parameters *-dosaf 1 -GL 1 -doCounts 1 -minmapq 20 -minq 20 -uniqueonly 1 -remove_bads 1*. All genomes were downsampled to similar coverages (6.5-7.5x) using *samtools v.1.9* [66] *view -s,* prior to heterozygosity estimation. For the identification of ROH segments, we used *PLINK v.1.9* [83] with parameters “*--homozyg-window-snp 50 --homozyg-window-het 1 --homozyg-snp 30 --homozyg-kb 500 --homozyg-density 30*”. We calculated F_ROH_, the proportion of the genome containing ROH segments, as the sum of ROH segments divided by the total size of the sheep reference genome. We grouped the ROHs into different size classes. In order to estimate the time period of inbreeding corresponding to each size class, we used the formula *g = 100/2rL* [38,39], where *g* corresponds to generation time, *r* to recombination rate and *L* to ROH length in Mb. We also estimated inbreeding time using the mean ROH length in CYM and ANM genomes. We used 1.5 cM/Mb as the recombination rate and calculated the estimated times using a generation time of 3 years.

### Mutation load and GERP scores

We downloaded GERP scores from the Ensembl database, which were calculated for domestic sheep reference Oar_v3.1 [84]. Using the UCSC liftover tool, we mapped the conservation scores to positions on the reference Oar_v4 [85]. The ancestral states were determined based on the alleles observed in the goat genome. We calculated the mutation load for each derived allele in sheep genomes in conserved regions of the genome, following von Seth and colleagues [41]. For this, we first defined conserved genomic regions in the reference genome as strings of consecutive bases with GERP>4. We then calculated the relative mutation load (RML) [41] for each genome using:

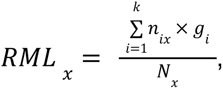

where *x* is one sheep genome, *k* corresponds to the total number of conserved regions, *g* is the GERP score for each region *i*, *n* is the number of derived alleles in region *i* in genome *x*, and *N* corresponds to the total number of derived alleles in genome *x*. The number of derived alleles was counted as one for heterozygous sites and as two for homozygous sites.

## Acknowledgements

We thank all the members of METU CompEvo and Hacettepe Human_G group, as well as Germán Hernández Alonso, M. Çelik, T. Hatipoğlu and H. Emir for their helpful suggestions and discussions.

## Data Availability

All newly generated sequence data were submitted to the European Nucleotide Archive (ENA) with the project ID PRJEB69690.

## References

1. Ceballos G, Ehrlich PR, Barnosky AD, García A, Pringle RM, Palmer TM. Accelerated modern human–induced species losses: Entering the sixth mass extinction. Sci Adv. 2015;1: e1400253. doi:10.1126/sciadv.1400253

2. Díez-del-Molino D, Sánchez-Barreiro F, Barnes I, Gilbert MTP, Dalén L. Quantifying Temporal Genomic Erosion in Endangered Species. Trends Ecol Evol. 2018;33: 176–185. doi:10.1016/j.tree.2017.12.002

3. Schmidt C, Hoban S, Hunter M, Paz-Vinas I, Garroway CJ. Genetic diversity and IUCN Red List status. Conserv Biol. 2023;37: e14064. doi:10.1111/cobi.14064

4. Garner BA, Hoban S, Luikart G. IUCN Red List and the value of integrating genetics. Conserv Genet. 2020;21: 795–801. doi:10.1007/s10592-020-01301-6

5. Frankham R. Genetics and extinction. Biol Conserv. 2005;126: 131–140. doi:10.1016/j.biocon.2005.05.002

6. Nabutanyi P, Wittmann MJ. Models for Eco-Evolutionary Extinction Vortices under Balancing Selection. Am Nat. 2021;197: 336–350. doi:10.1086/712805

7. Bosse M, van Loon S. Challenges in quantifying genome erosion for conservation. Front Genet. 2022;13. Available: https://www.frontiersin.org/articles/10.3389/fgene.2022.960958

8. Bertorelle G, Raffini F, Bosse M, Bortoluzzi C, Iannucci A, Trucchi E, et al. Genetic load: genomic estimates and applications in non-model animals. Nat Rev Genet. 2022;23: 492–503. doi:10.1038/s41576-022-00448-x

9. Arıhan O, Bilgin CC. Population biology and conservation of the Turkish mouflon (Ovis gmelini anatolica Valenciennes, 1856). In: Nahlik A, Uloth W, editors. Proceedings of the Third International Symposium on Mouflon. Sopron , Hungary: Institute of Wildlife Management; 2001. pp. 31–36.

10. Hadjisterkotis E. The Cyprus mouflon, a threatened species in a biodiversity “hotspot” area. 2001.

11. Hadjisterkotis ES, Bider JR. Reproduction of Cyprus mouflon Ovis gmelini ophion in captivity and in the wild. Int Zoo Yearb. 1993;32: 125–131. doi:10.1111/j.1748-1090.1993.tb03524.x

12. Michel S, Ghoddousi A. IUCN Red List of Threatened Species: Ovis gmelini. IUCN Red List Threat Species. 2020 [cited 4 Oct 2023]. Available: https://www.iucnredlist.org/en

13. Özdirek L. Estimation of demography and seasonal habitat use patterns of anatolian mouflon (ovis gmelinii anatolica) in Konya Bozdag protection area using distance sampling. Master Thesis, Middle East Technical University. 2009. Available: https://open.metu.edu.tr/handle/11511/18923

14. Özüt D. Evaluation of the adaptation process of a reintroduced anatolian mouflon (ovis gmelinii anatolica) population through studying its demography and spatial ecology. Yeniden aşılan bir Anadolu Yaban Koyunu (ovis gmelinii anatolica) toplumunun demografisi ve uzamsal ekolojisi araştıralarak uyum sürecinin değerlendirilmesi. 2009 [cited 4 Oct 2023]. Available: https://open.metu.edu.tr/handle/11511/19452

15. Hadjisterkotis E, Mereu P, Masala P. A review of the nomenclatural spelling variation of the Armenian mouflon (Ovis gmelini gmelinii) and the Cyprian mouflon (O. g. ophion). 2nd ed. Book of Abstracts – 6th World Congress on Mountain Ungulates and 5th International Symposium on Mouflon. 2nd ed. Nicosia: Ministry of the Interior; 2016. pp. 48–50.

16. Garel M, Marchand P, Bourgoin G, Santiago-Moreno J, Portanier E, Piegert H, et al. Mouflon Ovis gmelini Blyth, 1841. In: Hackländer K, Zachos FE, editors. Handbook of the Mammals of Europe. Cham: Springer International Publishing; 2020. pp. 1–35. doi:10.1007/978-3-319-65038-8_34-1

17. Abell JT, Quade J, Duru G, Mentzer SM, Stiner MC, Uzdurum M, et al. Urine salts elucidate Early Neolithic animal management at Aşıklı Höyük, Turkey. Sci Adv. 2019;5: eaaw0038. doi:10.1126/sciadv.aaw0038

18. Zeder MA. Domestication and early agriculture in the Mediterranean Basin: Origins, diffusion, and impact. Proc Natl Acad Sci U S A. 2008;105: 11597–11604. doi:10.1073/pnas.0801317105

19. Demirci S, Koban Baştanlar E, Dağtaş ND, Pişkin E, Engin A, Özer F, et al. Mitochondrial DNA Diversity of Modern, Ancient and Wild Sheep (Ovis gmelinii anatolica) from Turkey: New Insights on the Evolutionary History of Sheep. Barendse W, editor. PLoS ONE. 2013;8: e81952. doi:10.1371/journal.pone.0081952

20. Her C, Rezaei H-R, Hughes S, Naderi S, Duffraisse M, Mashkour M, et al. Broad maternal geographic origin of domestic sheep in Anatolia and the Zagros. Anim Genet. 2022;53: 452–459. doi:10.1111/age.13191

21. Guerrini M, Forcina G, Panayides P, Lorenzini R, Garel M, Anayiotos P, et al. Molecular DNA identity of the mouflon of Cyprus (Ovis orientalis ophion, Bovidae): Near Eastern origin and divergence from Western Mediterranean conspecific populations. Syst Biodivers. 2015. doi:10.1080/14772000.2015.1046409

22. Sanna D, Barbato M, Hadjisterkotis E, Cossu P, Decandia L, Trova S, et al. The First Mitogenome of the Cyprus Mouflon (Ovis gmelini ophion): New Insights into the Phylogeny of the Genus Ovis. PLoS ONE. 2015;10: e0144257. doi:10.1371/journal.pone.0144257

23. Vigne J-D, Carrère I, Briois F, Guilaine J. The Early Process of Mammal Domestication in the Near East: New Evidence from the Pre-Neolithic and Pre-Pottery Neolithic in Cyprus. Curr Anthropol. 2011;52: S255–S271. doi:10.1086/659306

24. Vigne J-D, Zazzo A, Cucchi T, Briois F, Guilaine J. The transportation of mammals to Cyprus shed light on early voyaging and boats in the mediterranean sea. Eurasian Prehistory. 2014;10: 157–176.

25. Arıhan O. Population biology spatial distribution and grouping patterns of the Anatolian Mouflon ovis gmelinii anatolica Valenciennes 1856. Master Thesis, Middle East Technical University. 2000. Available: https://open.metu.edu.tr/handle/11511/2696

26. Hadjisterkotis E. The Cyprus mouflon Ovis gmelini ophion : management, conservation and evolution. McGill University; 1992 [cited 8 Oct 2023]. Available: https://escholarship.mcgill.ca/concern/theses/3j333470z

27. Kence A, Tarhan S. Anatolian Mouflon. Wild Sheep Goats Their Relat Status Surv Conserv Action Plan Caprinae. 1997; 137–138.

28. Nicolaou H, Hadjisterkotis E, Papasavvas K. Past and present distribution and abundance of the Cyprus mouflon. 3rd ed. Book of Abstracts – 6th World Congress on Mountain Ungulates and 5th International Symposium on Mouflon. 3rd ed. Nicosia: Ministry of the Interior; 2016. p. 103.

29. Turan N. Türkiye’nin Av ve Yaban Hayvanları, Memeliler. Ankara: Self Published; 1984.

30. Kapnisis K, Kassinis N, Papanikolopoulou V, Diakou A. Endoparasites in wild populations of Cyprus mouflon (Ovis gmelini ophion). Vet Parasitol Reg Stud Rep. 2022;34: 100767. doi:10.1016/j.vprsr.2022.100767

31. Chen Z-H, Xu Y-X, Xie X-L, Wang D-F, Aguilar-Gómez D, Liu G-J, et al. Whole-genome sequence analysis unveils different origins of European and Asiatic mouflon and domestication-related genes in sheep. Commun Biol. 2021;4: 1–15. doi:10.1038/s42003-021-02817-4

32. Deng J, Xie X-L, Wang D-F, Zhao C, Lv F-H, Li X, et al. Paternal Origins and Migratory Episodes of Domestic Sheep. Curr Biol. 2020;30: 4085–4095.e6. doi:10.1016/j.cub.2020.07.077

33. Kijas JW, Lenstra JA, Hayes B, Boitard S, Neto LRP, Cristobal MS, et al. Genome-Wide Analysis of the World’s Sheep Breeds Reveals High Levels of Historic Mixture and Strong Recent Selection. PLOS Biol. 2012;10: e1001258. doi:10.1371/journal.pbio.1001258

34. Li X, Yang J, Shen M, Xie X-L, Liu G-J, Xu Y-X, et al. Whole-genome resequencing of wild and domestic sheep identifies genes associated with morphological and agronomic traits. Nat Commun. 2020;11: 2815. doi:10.1038/s41467-020-16485-1

35. Meadows J, Cemal I, Karaca O, Gootwine E, Kijas J. Five Ovine Mitochondrial Lineages Identified From Sheep Breeds of the Near East. Genetics. 2007;175: 1371–9. doi:10.1534/genetics.106.068353

36. Nadachowska-Brzyska K, Burri R, Smeds L, Ellegren H. PSMC analysis of effective population sizes in molecular ecology and its application to black-and-white Ficedula flycatchers. Mol Ecol. 2016;25: 1058–1072. doi:10.1111/mec.13540

37. Morell Miranda P, Soares AER, Günther T. Demographic reconstruction of the Western sheep expansion from whole-genome sequences. G3 GenesGenomesGenetics. 2023; jkad199. doi:10.1093/g3journal/jkad199

38. Kardos M, Åkesson M, Fountain T, Flagstad Ø, Liberg O, Olason P, et al. Genomic consequences of intensive inbreeding in an isolated wolf population. Nat Ecol Evol. 2018;2: 124–131. doi:10.1038/s41559-017-0375-4

39. Thompson EA. Identity by Descent: Variation in Meiosis, Across Genomes, and in Populations. Genetics. 2013;194: 301–326. doi:10.1534/genetics.112.148825

40. Cooper GM, Stone EA, Asimenos G, Green ED, Batzoglou S, Sidow A. Distribution and intensity of constraint in mammalian genomic sequence. Genome Res. 2005;15: 901–913. doi:10.1101/gr.3577405

41. von Seth J, Dussex N, Díez-del-Molino D, van der Valk T, Kutschera VE, Kierczak M, et al. Genomic insights into the conservation status of the world’s last remaining Sumatran rhinoceros populations. Nat Commun. 2021;12: 2393. doi:10.1038/s41467-021-22386-8

42. Chessa B, Pereira F, Arnaud F, Amorim A, Goyache F, Mainland I, et al. Revealing the History of Sheep Domestication Using Retrovirus Integrations. Science. 2009;324: 532–536. doi:10.1126/science.1170587

43. Hiendleder S, Kaupe B, Wassmuth R, Janke A. Molecular analysis of wild and domestic sheep questions current nomenclature and provides evidence for domestication from two different subspecies. Proc R Soc B Biol Sci. 2002;269: 893–904. doi:10.1098/rspb.2002.1975

44. Wang D-F, terWengel PO, Li M-H, Lv F-H. Genomic analyses of Asiatic Mouflon in Iran provide insights into the domestication and evolution of sheep. bioRxiv; 2023. p. 2023.10.06.561316. doi:10.1101/2023.10.06.561316

45. Cheng H, Zhang Z, Wen J, Lenstra JA, Heller R, Cai Y, et al. Long divergent haplotypes introgressed from wild sheep are associated with distinct morphological and adaptive characteristics in domestic sheep. PLoS Genet. 2023;19: e1010615. doi:10.1371/journal.pgen.1010615

46. Yurtman E, Özer O, Yüncü E, Dağtaş ND, Koptekin D, Çakan YG, et al. Archaeogenetic analysis of Neolithic sheep from Anatolia suggests a complex demographic history since domestication. Commun Biol. 2021;4: 1–11. doi:10.1038/s42003-021-02794-8

47. Hadjisterkotis E. Seasonal and monthly distribution of deaths of Cyprus Mouflon *Ovis gmelini ophion*. Pirineos. 2002;157: 81–88. doi:10.3989/pirineos.2002.v157.63

48. Hadjisterkotis E, van Haaften JL. Die Niederwildjagd im Wald von Paphos und ihre Auswirkungen auf das gefährdete zyprische MufflonOvis gmelini ophion. Z Für Jagdwiss. 1997;43: 279–282. doi:10.1007/BF02239894

49. Toumazos P, Hadjisterkotis E. Diseases of the Cyprus mouflon as determined by Standard gross and histopathological methods. Proceedings of the Second International Symposium on Mediterranean Mouflon. Nicosia: Game Fund; 1997. pp. 150–161.

50. Mathur S, Tomeček JM, Tarango-Arámbula LA, Perez RM, DeWoody JA. An evolutionary perspective on genetic load in small, isolated populations as informed by whole genome resequencing and forward-time simulations. Evolution. 2023;77: 690–704. doi:10.1093/evolut/qpac061

51. Robinson J, Kyriazis CC, Yuan SC, Lohmueller KE. Deleterious Variation in Natural Populations and Implications for Conservation Genetics. Annu Rev Anim Biosci. 2023;11: 93–114. doi:10.1146/annurev-animal-080522-093311

52. Carpenter ML, Buenrostro JD, Valdiosera C, Schroeder H, Allentoft ME, Sikora M, et al. Pulling out the 1%: Whole-Genome Capture for the Targeted Enrichment of Ancient DNA Sequencing Libraries. Am J Hum Genet. 2013;93: 852–864. doi:10.1016/j.ajhg.2013.10.002

53. Linnell JDC, Zachos FE. Status and distribution patterns of European ungulates: genetics, population history and conservation. In: Apollonio M, Andersen R, Putman R, editors. Ungulate Management in Europe: Problems and Practices. Cambridge: Cambridge University Press; 2011. pp. 12–53. doi:10.1017/CBO9780511974137.003

54. Hare MP, Nunney L, Schwartz MK, Ruzzante DE, Burford M, Waples RS, et al. Understanding and Estimating Effective Population Size for Practical Application in Marine Species Management. Conserv Biol. 2011;25: 438–449. doi:10.1111/j.1523-1739.2010.01637.x

55. Cairns KM. What is a dingo – origins, hybridisation and identity. Aust Zool. 2021;41: 322–337. doi:10.7882/AZ.2021.004

56. Cairns KM, Brown SK, Sacks BN, Ballard JWO. Conservation implications for dingoes from the maternal and paternal genome: Multiple populations, dog introgression, and demography. Ecol Evol. 2017;7: 9787–9807. doi:10.1002/ece3.3487

57. Elledge AE, Leung LK-P, Allen LR, Firestone K, Wilton AN. Assessing the taxonomic status of dingoes Canis familiaris dingo for conservation. Mammal Rev. 2006;36: 142–156. doi:10.1111/j.1365-2907.2006.00086.x

58. Jhala Y, Boitani L, Phillips M. IUCN Red List of Threatened Species: Canis lupus. IUCN Red List Threat Species. 2018 [cited 5 Nov 2023]. Available: https://www.iucnredlist.org/en

59. Barbato M, Masseti M, Pirastru M, Columbano N, Scali M, Vignani R, et al. Islands as Time Capsules for Genetic Diversity Conservation: The Case of the Giglio Island Mouflon. Diversity. 2022;14: 609. doi:10.3390/d14080609

60. Mereu P, Pirastru M, Barbato M, Satta V, Hadjisterkotis E, Manca L, et al. Identification of an ancestral haplotype in the mitochondrial phylogeny of the ovine haplogroup B. PeerJ. 2019;7: e7895. doi:10.7717/peerj.7895

61. Portanier E, Chevret P, Gélin P, Benedetti P, Sanchis F, Barbanera F, et al. New insights into the past and recent evolutionary history of the Corsican mouflon (Ovis gmelini musimon) to inform its conservation. Conserv Genet. 2022;23: 91–107. doi:10.1007/s10592-021-01399-2

62. Satta V, Mereu P, Barbato M, Pirastru M, Bassu G, Manca L, et al. Genetic characterization and implications for conservation of the last autochthonous Mouflon population in Europe. Sci Rep. 2021;11: 14729. doi:10.1038/s41598-021-94134-3

63. Meyer M, Kircher M. Illumina sequencing library preparation for highly multiplexed target capture and sequencing. Cold Spring Harb Protoc. 2010;2010: pdb.prot5448. doi:10.1101/pdb.prot5448

64. Schubert M, Lindgreen S, Orlando L. AdapterRemoval v2: rapid adapter trimming, identification, and read merging. BMC Res Notes. 2016;9: 88. doi:10.1186/s13104-016-1900-2

65. Li H, Durbin R. Fast and accurate long-read alignment with Burrows–Wheeler transform. Bioinformatics. 2010;26: 589–595. doi:10.1093/bioinformatics/btp698

66. Li H, Handsaker B, Wysoker A, Fennell T, Ruan J, Homer N, et al. The Sequence Alignment/Map format and SAMtools. Bioinformatics. 2009;25: 2078–2079. doi:10.1093/bioinformatics/btp352

67. Poplin R, Ruano-Rubio V, DePristo MA, Fennell TJ, Carneiro MO, Auwera GAV der, et al. Scaling accurate genetic variant discovery to tens of thousands of samples. bioRxiv; 2018. p. 201178. doi:10.1101/201178

68. Li H. A statistical framework for SNP calling, mutation discovery, association mapping and population genetical parameter estimation from sequencing data. Bioinformatics. 2011;27: 2987–2993. doi:10.1093/bioinformatics/btr509

69. Mittnik A, Wang C-C, Svoboda J, Krause J. A Molecular Approach to the Sexing of the Triple Burial at the Upper Paleolithic Site of Dolní Věstonice. PLOS ONE. 2016;11: e0163019. doi:10.1371/journal.pone.0163019

70. Kuhn JMM, Jakobsson M, Günther T. Estimating genetic kin relationships in prehistoric populations. PLOS ONE. 2018;13: e0195491. doi:10.1371/journal.pone.0195491

71. Korneliussen TS, Albrechtsen A, Nielsen R. ANGSD: Analysis of Next Generation Sequencing Data. BMC Bioinformatics. 2014;15: 356. doi:10.1186/s12859-014-0356-4

72. Katoh K, Standley DM. MAFFT Multiple Sequence Alignment Software Version 7: Improvements in Performance and Usability. Mol Biol Evol. 2013;30: 772–780. doi:10.1093/molbev/mst010

73. Bandelt HJ, Forster P, Röhl A. Median-joining networks for inferring intraspecific phylogenies. Mol Biol Evol. 1999;16: 37–48. doi:10.1093/oxfordjournals.molbev.a026036

74. Leigh JW, Bryant D. popart: full-feature software for haplotype network construction. Methods Ecol Evol. 2015;6: 1110–1116. doi:10.1111/2041-210X.12410

75. Maier, Patterson. admixtools: Tools for inferring demographic history from genetic data in uqrmaie1/admixtools: Inferring demographic history from genetic data. 2023. Available: https://rdrr.io/github/uqrmaie1/admixtools/man/admixtools.html

76. Patterson N, Moorjani P, Luo Y, Mallick S, Rohland N, Zhan Y, et al. Ancient Admixture in Human History. Genetics. 2012;192: 1065–1093. doi:10.1534/genetics.112.145037

77. Koptekin D, Yüncü E, Rodríguez-Varela R, Altınışık NE, Psonis N, Kashuba N, et al. Spatial and temporal heterogeneity in human mobility patterns in Holocene Southwest Asia and the East Mediterranean. Curr Biol. 2023;33: 41–57.e15. doi:10.1016/j.cub.2022.11.034

78. Paradis E, Schliep K. ape 5.0: an environment for modern phylogenetics and evolutionary analyses in R. Bioinformatics. 2019;35: 526–528. doi:10.1093/bioinformatics/bty633

79. Li H, Durbin R. Inference of Human Population History From Whole Genome Sequence of A Single Individual. Nature. 2011;475: 493–496. doi:10.1038/nature10231

80. Chen L, Qiu Q, Jiang Y, Wang K, Lin Z, Li Z, et al. Large-scale ruminant genome sequencing provides insights into their evolution and distinct traits. Science. 2019;364: eaav6202. doi:10.1126/science.aav6202

81. Lv F-H, Cao Y-H, Liu G-J, Luo L-Y, Lu R, Liu M-J, et al. Whole-Genome Resequencing of Worldwide Wild and Domestic Sheep Elucidates Genetic Diversity, Introgression, and Agronomically Important Loci. Mol Biol Evol. 2021;39: msab353. doi:10.1093/molbev/msab353

82. Zhao Y-X, Yang J, Lv F-H, Hu X-J, Xie X-L, Zhang M, et al. Genomic Reconstruction of the History of Native Sheep Reveals the Peopling Patterns of Nomads and the Expansion of Early Pastoralism in East Asia. Mol Biol Evol. 2017;34: 2380–2395. doi:10.1093/molbev/msx181

83. Chang CC, Chow CC, Tellier LC, Vattikuti S, Purcell SM, Lee JJ. Second-generation PLINK: rising to the challenge of larger and richer datasets. GigaScience. 2015;4: s13742-015-0047–8. doi:10.1186/s13742-015-0047-8

84. Martin FJ, Amode MR, Aneja A, Austine-Orimoloye O, Azov AG, Barnes I, et al. Ensembl 2023. Nucleic Acids Res. 2023;51: D933–D941. doi:10.1093/nar/gkac958

85. Nassar LR, Barber GP, Benet-Pagès A, Casper J, Clawson H, Diekhans M, et al. The UCSC Genome Browser database: 2023 update. Nucleic Acids Res. 2023;51: D1188–D1195. doi:10.1093/nar/gkac1072

