## Supplemental Figures for "Population genomic history of the endangered Anatolian and Cyprian mouflons in relation to worldwide wild, feral and domestic sheep lineages"

### Supplementary Figures

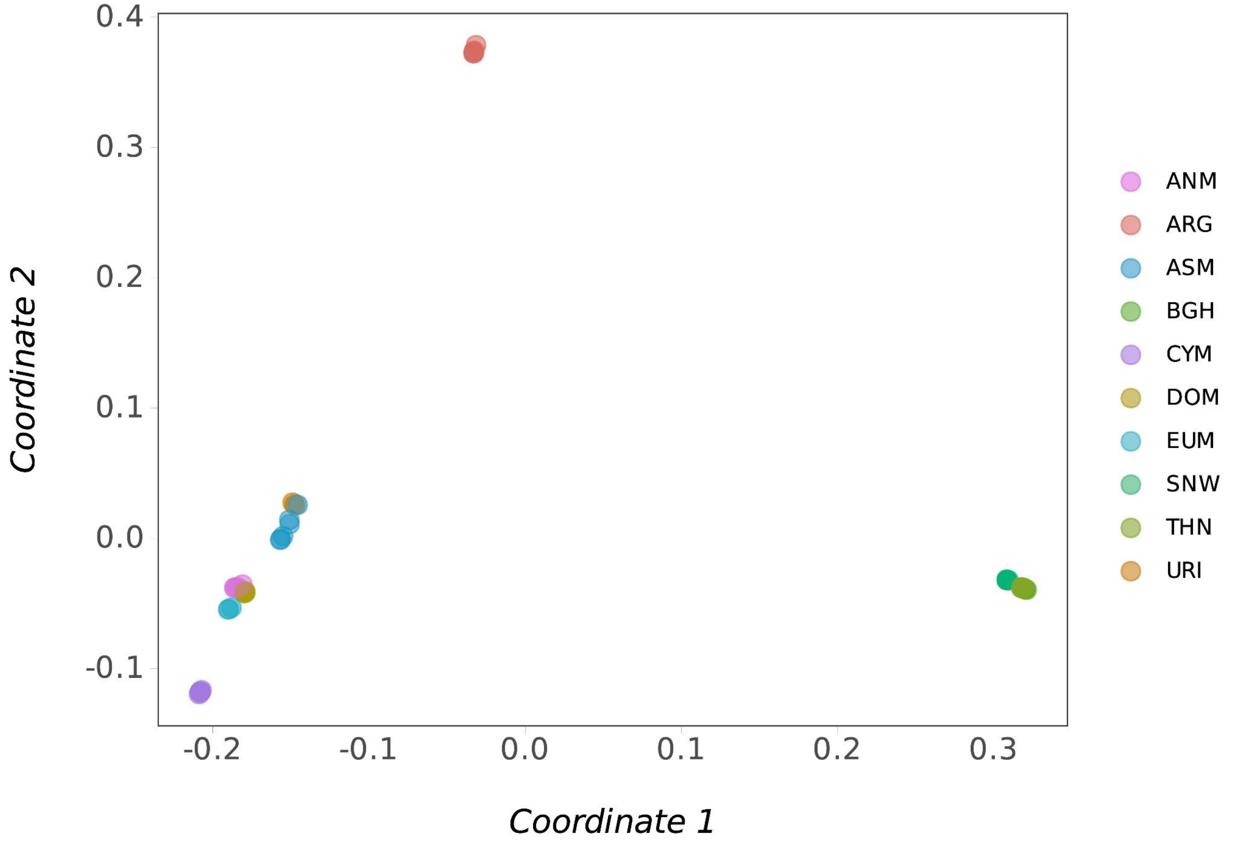


**Figure S1. MDS analysis of the studied individuals.** Multidimensional scaling using 1 - *outgroup-$f_3$* values between individuals as distance proxies, with goat as the outgroup. Coordinate 1 versus 2 is plotted.

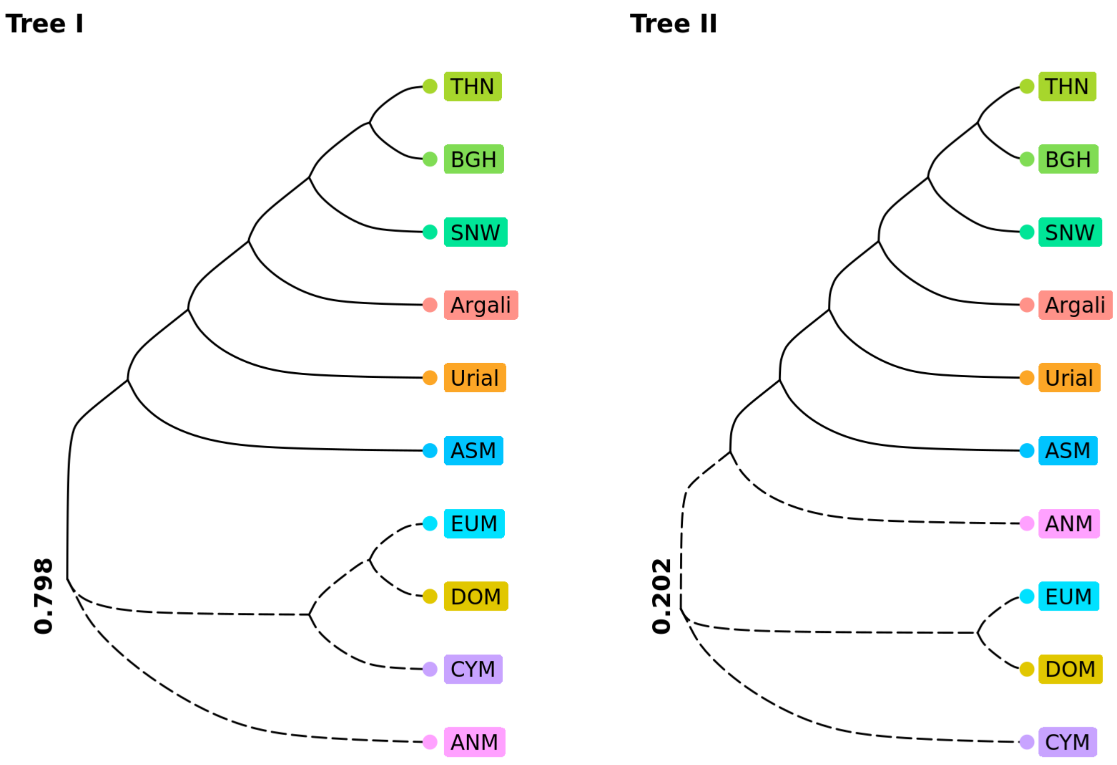


**Figure S2. Proportion of NJ trees observed amongst bootstraps.** Neighbour-joining trees constructed using 1 - *outgroup-*$f_3$ statistics as genetic distances. Sampling chunks of SNPs with replacement, 500 bootstraps were performed in total. Tree I was observed in 80% of the bootstraps, while Tree II was observed 20% of the time.

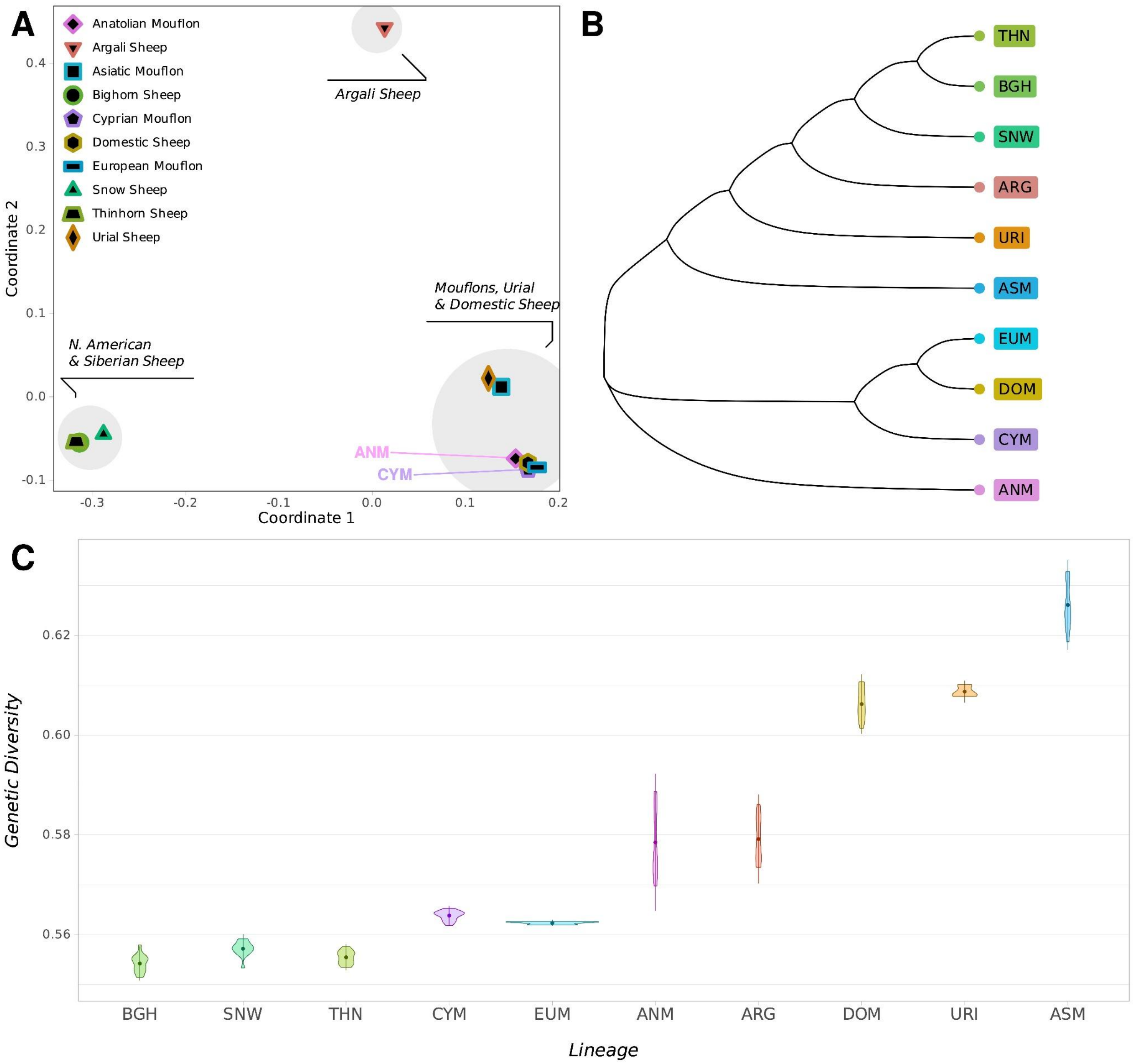


**Figure S3. Phylogenetic relationships between these sheep lineages and diversity estimates using the dataset of goat heterozygous SNPs.** (A) Multidimensional scaling (MDS) analysis of the studied sheep lineages, using 1-outgroup $f_3$ statistics as distance proxies. (B) Neighbour-joining (NJ) tree of the studied sheep lineages, using (1 - outgroup $f_3$) as distance proxies. (C) Within-population diversity values estimated using pairwise 1 - outgroup f3 statistics per lineage.

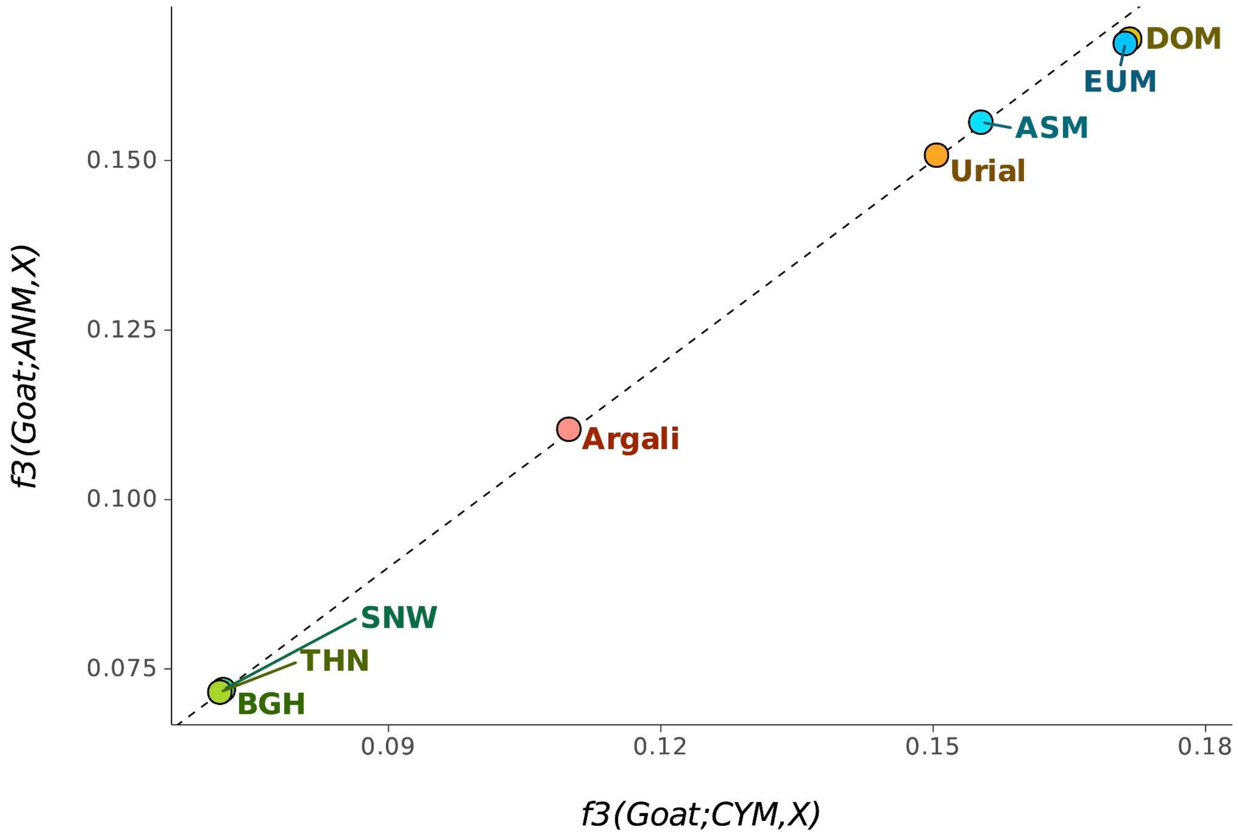


**Figure S4. Comparison of $f_3$ statistics between CYM/ANM and other sheep lineages.** *Outgroup-$f_3$* statistics of the form $f_3(Goat;CYM,X)$ on the x-axis plotted against $f_3(Goat;ANM,X)$ on the y-axis where X corresponds to other studied sheep lineages. Points on the dotted line represent similar affinity of both CYM and ANM to the other lineages.

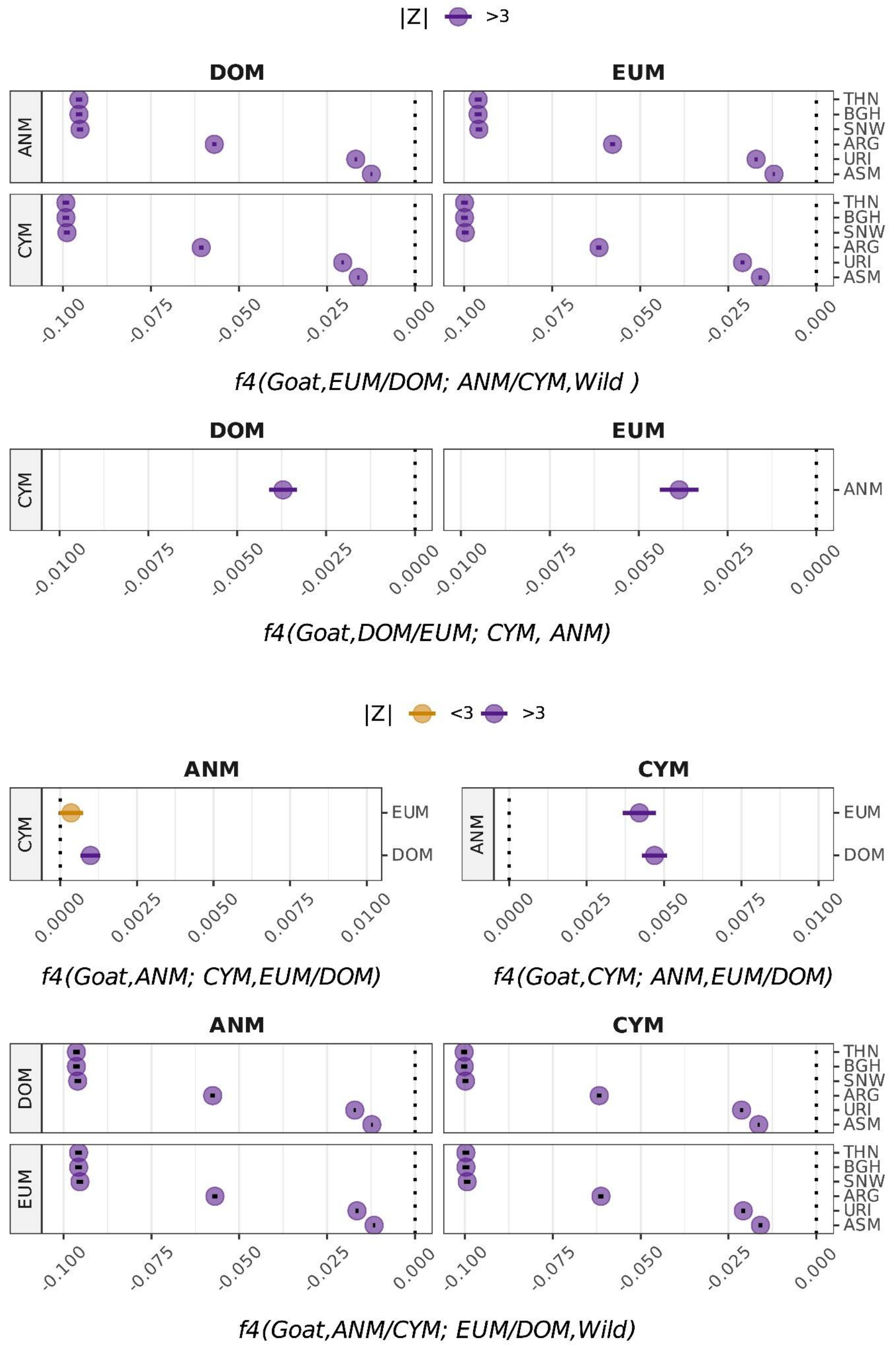


**Figure S5. $f_4$ statistics between CYM, ANM, DOM and EUM.** f4 statistics of the form *$f_4$(Goat, EUM/DOM, ANM/CYM, Wild)*, *$f_4$(Goat, EUM/DOM; ANM,CYM)*, *$f_4$(Goat, ANM/CYM; ANM/CYM, EUM/DOM)* and *$f_4$(Goat, ANM/CYM,; EUM/DOM,Wild)*, where

Wild corresponds to other studied sheep lineages. Orange color depicts results with $|Z| < 3$, and purple depicts significant results with $|Z| > 3$.

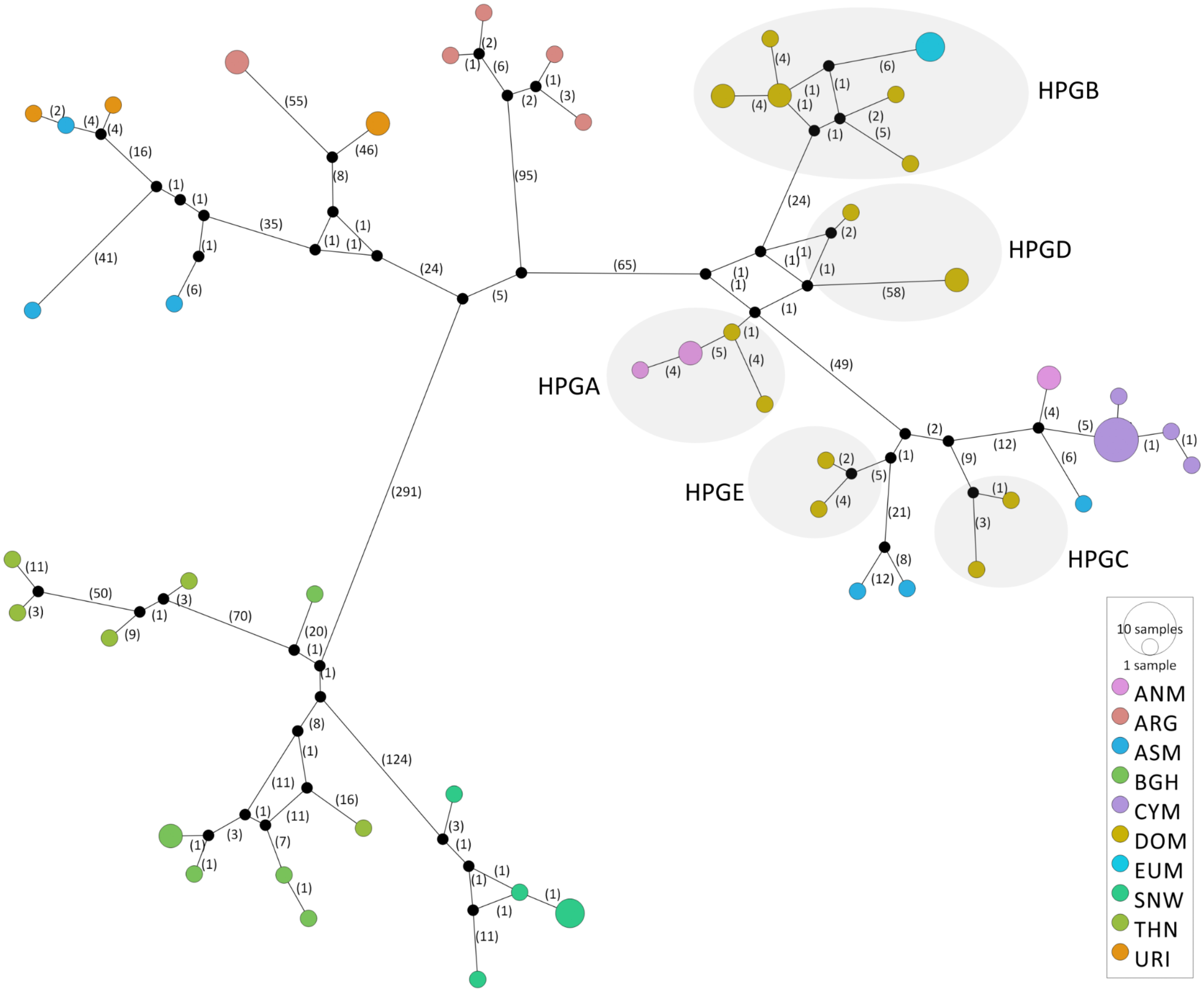


**Figure S6. Mitogenome DNA median-joining (MJ) network of wild and domestic sheep.** Node sizes are proportional to the number of samples in the node and numbers on edges show the number of nucleotide differences between nodes. Domestic haplogroup clusters were denoted by gray circles. Black dots represent hypothetical nodes.

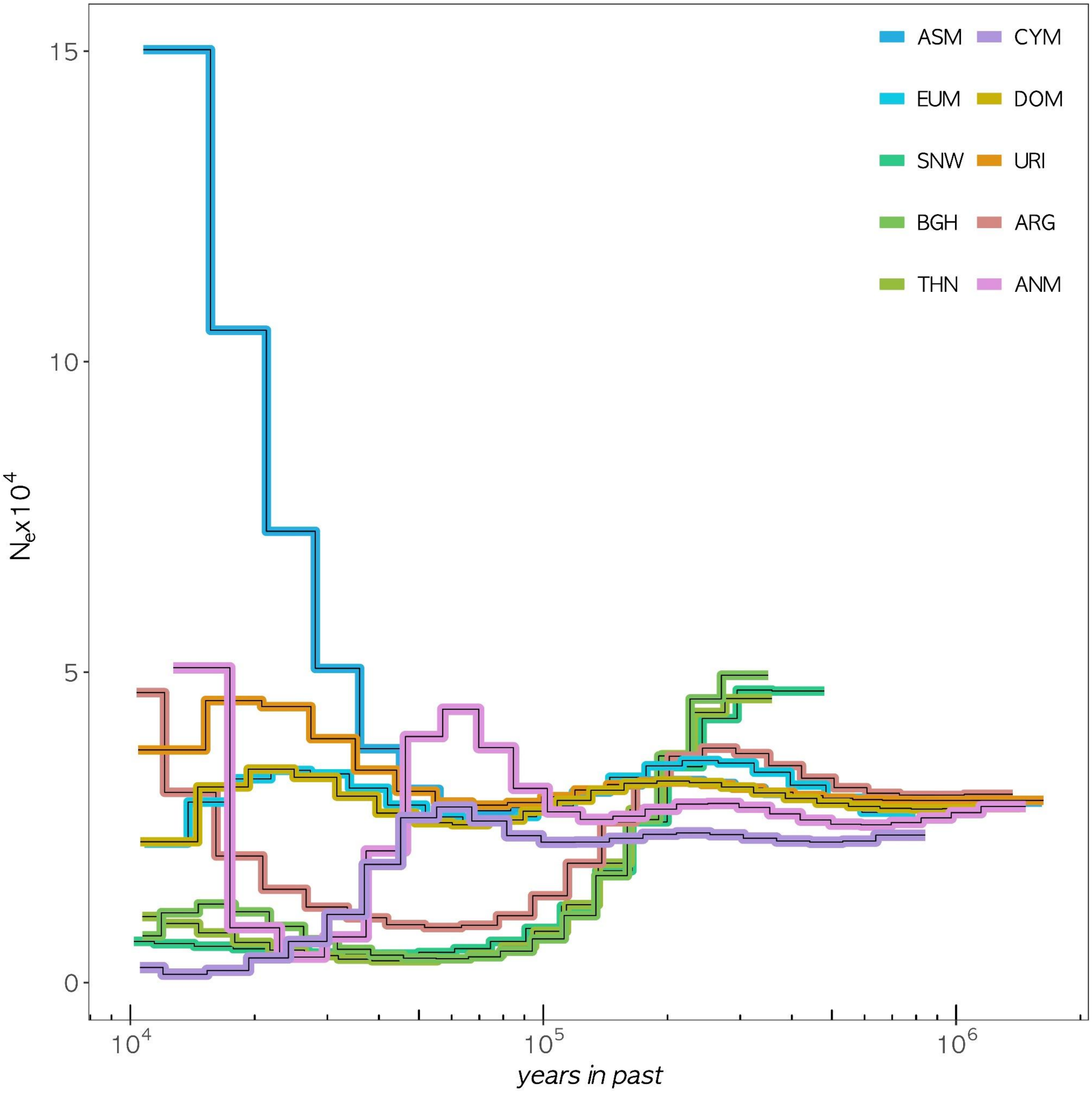


**Figure S7. PSMC analysis of genomes downsampled to similar coverages.** The genomes were downsampled to coverages 7.5-8.5x. Analysis was carried out assuming a generation time of 3 years and a mutation rate of $1.5x10^{-8}$. The x-axis shows time in a log scale, the y-axis shows the estimated effective population size.

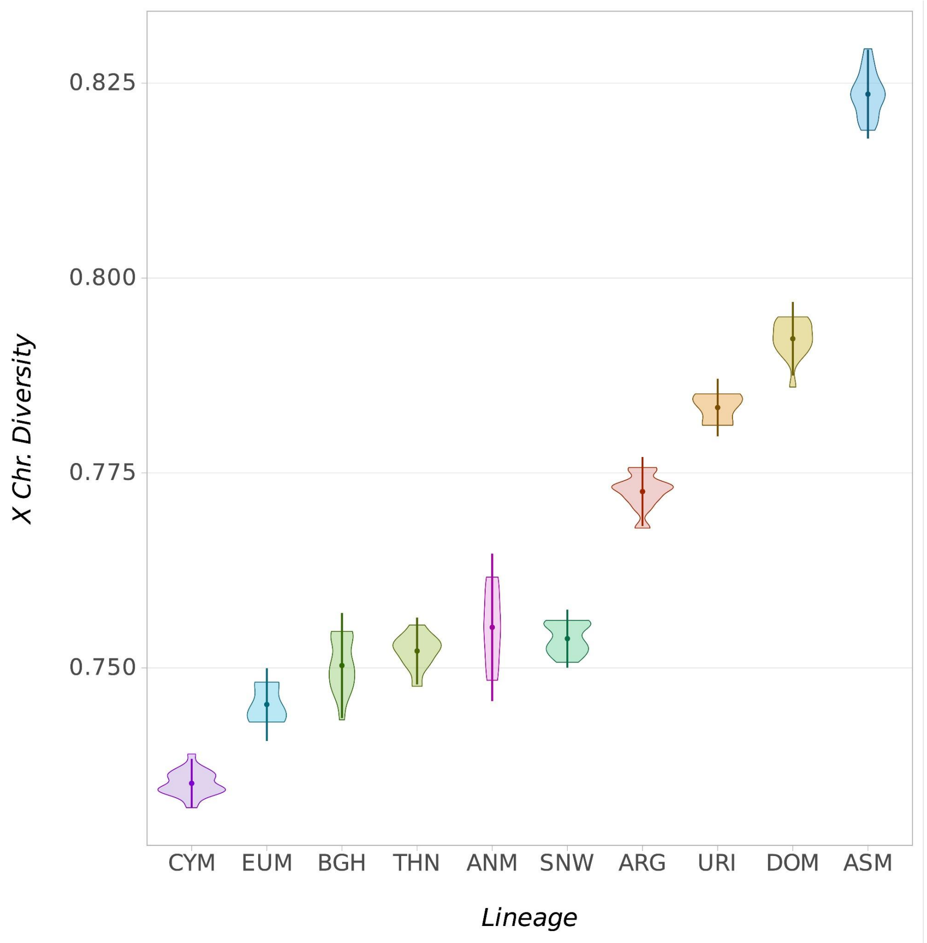


**Figure S8. X chromosome diversities of the studied sheep lineages.** Within-population X chr. diversity values estimated using pairwise 1 - outgroup-f3 statistics per lineage.

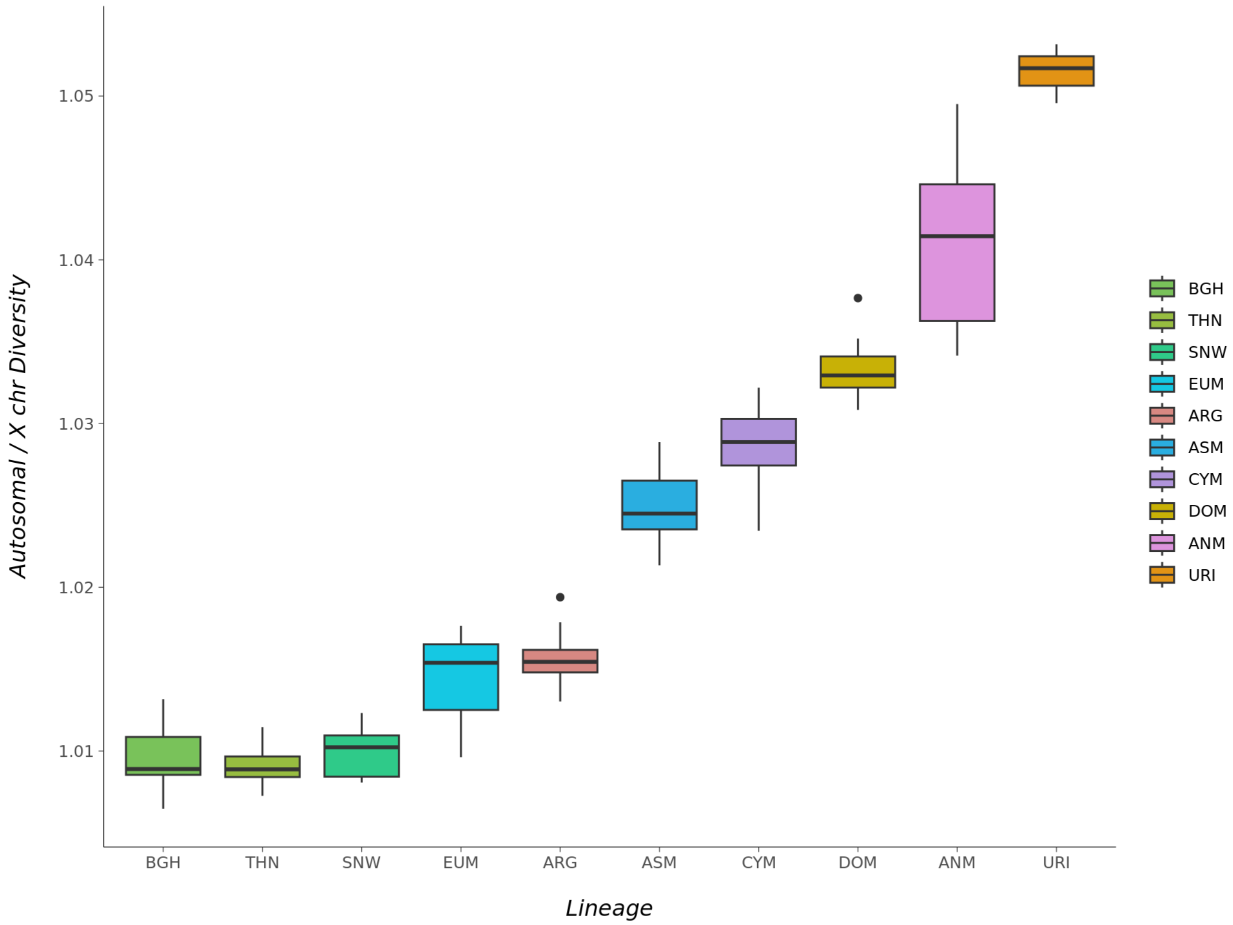


**Figure S9. Proportion of autosomal / X chromosome diversities of the studied sheep lineages.** Within-population autosomal and X chr. diversity values estimated using pairwise 1 - outgroup-f3 statistics per lineage.

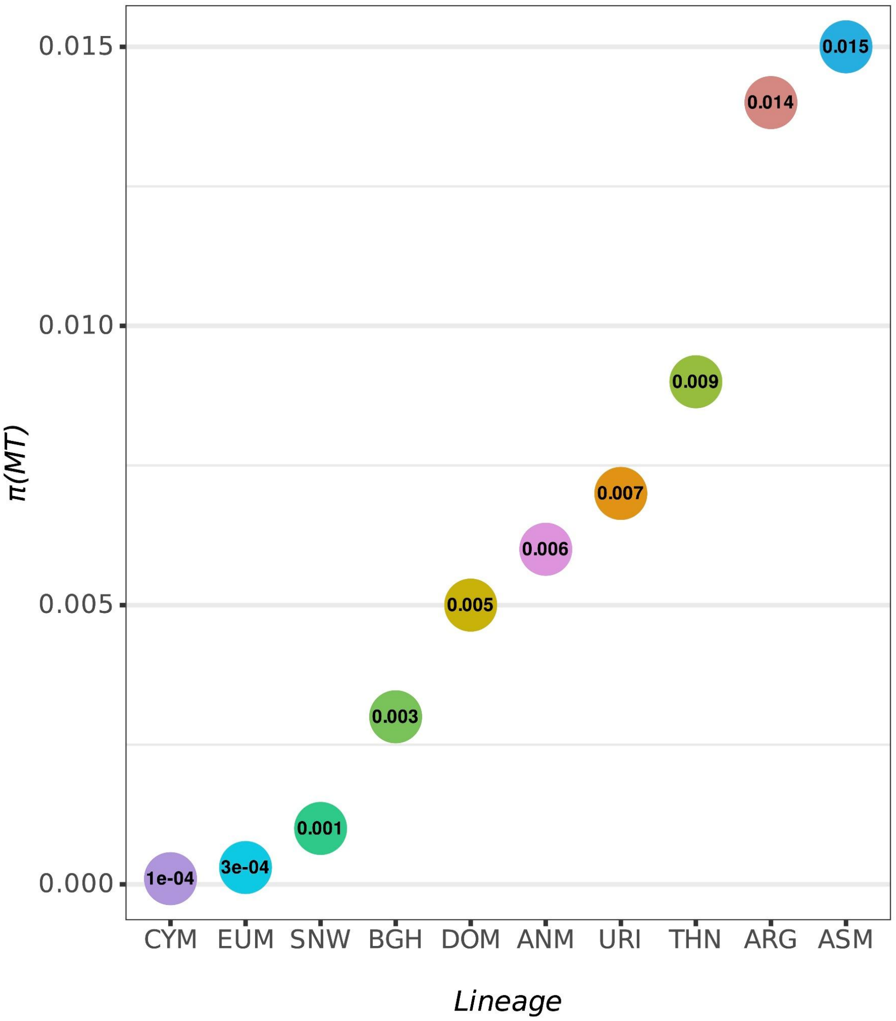


**Figure S10. Mitochondrial diversities of the studied sheep lineages.** Within population pi (π) estimates using mitochondrial genomes.
