## Supplemental Note for "Population genomic history of the endangered Anatolian and Cyprian mouflons in relation to worldwide wild, feral and domestic sheep lineages"

#### Supplemental Note 1:

### The Cyprus mouflon (*Ovis gmelini ophion*): a short historic review of the population numbers and the factors affecting its past and present decline

by Eleftherios Hadjisterkotis

##### Introduction

The Cyprian mouflon is an endangered species mainly inhabiting Pafos forest in Cyprus, within an area of 620 km$^2$ (Hadjisterkotis, 1993; 1999; Tsintides et al., 2016). There is only one population of this subspecies in a single reserve which is threatened by habitat degradation, forest fires, diseases transmitted by domestic sheep and goats foraging in the same area as mouflon (Hadjisterkotis, 1999), stray dogs (Hadjisterkotis and Bider, 1992), and particularly poaching (Hadjisterkotis, 1993; Hadjisterkotis and van Haaftern, 1997). Because of persistent poaching, several times the mouflons have come close to extinction (Hadjisterkotis and van Haaften, 1997). Although during the last decade in the south-east section of its habitat the population expanded its range towards the Troodos National Park, it remains on the list of the IUCN endangered species (Nicolaou et al., 2016). Regardless of this expansion, in the north-western part of its range the population is declining, which may have been a reason for the relatively high homozygosity observed. In order to understand the causes that brought the Cyprian mouflon several times close to extinction, as well as the recent drastic population reduction in the northern part of its range, the aim of this study is to provide information on the population numbers of the Cyprian mouflon through the ages, with emphasis on the most recent developments on management and conservation, which might have contributed to the observed high homozygosity in the northern part of its range.

##### Methods and Materials

Data were collected from a literature review, by examining the files of the Department of Forestry at Stavros tis Psokas forest station, personal observations and information from the personnel of the Department of Forestry.

##### Results and Discussion

The first records of mouflon on the island of Cyprus derived from a Neolithic settlement near the modern village of Parekklisha – in a locality known as *Shillourokampos*. According to detailed archaeological data (Guilaine et al., 1995; Guilaine, 2003; Guilaine and Briois, 2007), the long period of settlement at Shillourokampos can be divided into four main phases from 8.400 to 7.000 B.C. During the period of 8.400 – 8.100 B.C. there is evidence of wheat cultivation and legumes (Vinge et al., 2012) and the presence of a small sized wild boar (Vigne et al,. 2009). After a brief abandonment of the site, or a gap due to erosion in the period of 7.900-7.600 B.C. archaeological finding revealed a reduction in the number of wild boar, and for the first time there were mouflons (*Ovis gmelini*) (Vigne et al., 2014; Gerolemou and Hadjisterkotis, 2016). During the period of

7.600-7.400 B.C. there was a reduction in sheep bones suggesting a collapse or a failure in sheep production (Vigne et al. 2014). This collapse forced the inhabitants of *Shillourokampos* to introduce new sheep breeds for meat production, perhaps from the mainland, or from other unknown Cypriot Neolithic sites, or perhaps from feral sheep or wild mouflon present on the mountains, not yet found in any fossil sites (Garel, 2020; Sanna et al., 2015).

Records of Cyprus mouflon through the early Greek (2000 B.C. – 57 B.C.) and the Roman times (58 B.C. – 330 A.D.) are only reflected in art. However, the following account of the history of the mouflon from the Middle Ages to 1969 has been taken from the accounts of J. Biddulph, High Commissioner of Cyprus (1884), and the files and/or undated pamphlets obtained from the Department of Forestry of Cyprus compiled by Hadjisterkotis (1992). Although the accuracy and precision of the reported population estimates are impossible to evaluate, the general tendencies are probably reflected and thus used to outline the history.

In the Middle Ages, the mouflon was plentiful in nearly all parts of Cyprus and was hunted by the aristocracy using hounds and coursing them with cheetahs. By 1878 the mouflon were found in the Troodos Mountains in the western portion of the island (Hadjisterkotis, 1992, Fig. 1.1. page 15), where they were disturbed only by occasional woodcutters and peasants herding goats and sheep. Biddulph (1884) claims that at the time of the British occupation mouflon had been exterminated with the exception of a single flock of 25 members and a ban was placed on their hunting. By 1884 their numbers were thought to be increasing. In 1930 there was a flock of 20 or more in the Troodos forest and a similar number in the Pafos forest. By 1937 the population of the Pafos forest was thought to have been reduced to 15 animals through persisted poaching (Hadjisterkotis, 1992).

To protect the mouflon from poaching, on the 4$^{th}$ of November 1938 the entire Pafos forest (620 sq. Km.) was declared a permanent hunting preserve area, thus eliminating all forms of hunting from the area. This included all private properties of the three villages Kampos, Tsakistra and Mylikouri, and the Holy Royal Monastery of Saint Mary of Kykkos, found inside the forest (see map in Hadjisterkotis and van Haaften, 1997). In addition, the same year the Game Law was amended and strengthened to provide better protection for mouflon. Special forest staff and police were assigned full-time in order to reduce poaching. From 1938 to 1940 all goats were removed from the forest to eliminate competition with mouflon. Furthermore, it eliminated from the forest the goatherders, who were thought to be extensively involved with poaching. With a new decree on 16.11.1944 the above private properties surrounded by Pafos forest were excluded from the game preserve.

In 1950, a captive breeding stock was established at Stavros tis Psokas Forest Station with one male and two females. In 1955 a similar stock was established at the Limassol Zoological Garden from animals that originated from the captive stock from Stavros tis Psokas (Hadjisterkotis, 1993; Hadjisterkotis & Bider, 1993; Hadjisterkotis & Lambrou, 2001).

In 1970, Dr Van Haaften (1974), estimated the population to be 200 animals. He believed that there was no food shortage, and he recommended the planting of lure crops near the forest to protect agricultural crops from mouflon depredation. Until this time period, the population was restricted inside Pafos forest, due to extensive poaching by the local people living in villages around Paphos forest. People hunted mouflon for its meat, but also to keep them away from their vineyards and the rest of their crops. Following the Turkish invasion of Cyprus in 1974 and the removal of the Greek population from the Northern part of the island to the southern part, and the transfer of the Turkish Cypriot population to the North, the inhabitants of five Turkish Cypriot villages bordering Pafos forest (Vretsia, Anadiou, Sarama, Kios, and Pelathousa) moved to the Northern part of the island, leaving their fields and vineyards free for the mouflon to expand outside the border of the forest, into new agricultural areas reach in forage. In 1985, when I began my field work for my Ph.D. thesis dissertation in this area, there were several flocks of mouflon lambing in the nearby cliffs near the edge of the forest, and foraging in the forest-agricultural area of these villages (Hadjisterkotis, 1992). However, during a population survey that I did in 2010, the number of mouflon in this area was negligible. As I was informed from the President of the Community Council of the nearby village of Kannaviou, the cause of this drastic reduction was poaching. The same year, a senior officer from the Game and Fauna Service, in a newspaper interview dated 13th of September 2010, stated that small groups of poachers are making their living from mouflon poaching, earning tens of thousands of euros every year (Evripidou, 2010).

In 1983 the Department of Forestry estimated the mouflon population to be 500-600 animals. By 1988, the total estimate was approximately 2,000 mouflon. These estimates were based on the animals seen by employees of Stavros tis Psokas forest station when traveling the roads.

However, helicopter censuses flown in February 1992, under the direction of Prof. Roger Bider from McGill University, in cooperation with E. Hadjisterkotis, estimated the population to be 900-1500 animals. Thirty percent of the population were living inside Pafos forest and 70% were found at the margin of the forest. This census suggested that numbers in the forest have declined since 1988, perhaps as much as 50%, while those at the forest edge may have remained stable (Hadjisterkotis, 1992).

In 1993 the Game and Fauna Service (Game Fund) for the first time in its history assigned under my direction two Game Wardens for the management and conservation of mouflon, until my resignation from the Service in 1996. However, with only two Game Wardens was impossible to patrol and to protect the entire forest from poachers.

In 1996 a new development took place, which brought a major change in the protected status of the eastern part of Pafos forest. The Council of Ministers (upon the request of the people from the villages Kampos and Tsakistra and the approval of the Game Fund Committee), permitted small game hunting in the Limnitis valley and the nearby villages of Kampos and Tsakkistra, an area of about 40 $Km^2$. Hunting was permitted for chukar partridge (*Alectoris chukar Cypriotes*) and hare (*Lepus capensis*) every Wednesday and Sunday during November and December each year, with hunting dogs or without hunting dogs. Until this time, this type of hunting was permitted only outside the borders of Pafos forest. In addition, in the same hunting areas it is permitted to hunt woodpigeon and thrush, from November until the end of February Wednesdays and Sundays. Usually from the 20th until the 28th of February it is permitted on a daily basis

(Hadjisterkotis and Haaften 1997). This was followed by reports in the newspapers of shooting of up to 200 mouflons by poachers. These reports were investigated a few days later by the police and were considered groundless, since there was no evidence available to arrest anyone (Hadjisterkotis & van Haaftern, 1999).

As it was noted by Hadjisterkotis and van Haaftern (1997), since the authorities for game management did not provide the necessary manpower to prevent poaching of mouflon, this gave the opportunity to hunters and poachers to carry legally guns inside Paphos forest, and - although illegal - to shoot mouflon and/or to chase them with hunting dogs. During the hunting season, a number of “gun shy” hunting dogs run away from their owners, living in the wild as stray dogs. In addition, a number of hunters who are not satisfied with the hunting performance of their dogs, let them loose in the wild. These stray dogs form groups which, in Pafos forest, are chasing and killing mouflon and elsewhere might kill domestic sheep, sometimes causing serious losses to sheep herders. As I was informed from foresters working in the area of Kampos forest station, during the last few years there were high numbers of dogs, forming flocks of up to 15 animals. In several cases dogs were seen chasing mouflon, and it is not unusual to locate such wild sheep with cut throats. This phenomenon brought repeated complaints on public media mainly from foresters, complaining about insufficient protection of mouflon, and the observed drastic reduction in population numbers in the area of Kampos, the area in which our sample collection took place for this study.

Based on a long-term study of the seasonal and monthly distribution of deaths of the Cyprian mouflon from 1985 – 1998, two hundred and fifty-one mouflon carcasses were collected (Hadjisterkotis, 2002). One hundred and five (54 M, 37F, 13L, 1 adult of unknown sex) mouflon were identified as freshly dead, and underwent postmortem examination. The most important mortality factors were road kills, poaching, falling from cliffs, and predation from stray dogs. According to Toumazos and Hadjisterkotis (1997), poaching accounts for 30% of the mortality in mouflon, and roadkill 11%. If we consider roadkill as an intentional technique of poaching, then poaching accounts for 41% of the dead animals. Dead animals from stray dogs were found to be 13%.

A recent study (Kassinis et al., 2016), for the years 2011-2015 (99 specimens) showed disease-related mortality 30%, predation by feral dogs and foxes 25%, poaching 16% and collision with cars 13%, with the rest being accidents, snake bites and unknown. Mortality causes from 31 mouflon radio-tagged in years 2002-2007, indicated relatively higher levels of predation related mortality (32%) and poaching related (19%). Considering that, one in every three dead mouflon was killed by predators and one in five by poachers, over half of the animals found dead were killed by stray dogs and poachers. Comparing the above with Toumazos and Hadjisterkotis (1997) results (13% VS 32% mortality from stray dogs), it is obvious that in recent years the number mouflon found dead from stray dogs has almost tripled. Based on recent information from foresters from the Kampos Forestry Station, it seems that these stray dogs became permanent residents of Paphos forest, and are surviving in the wild by killing and consuming mouflon (I. Papadopoulos, personal communication, 2023). Considering that recently a number of intact animals were found with their carotids cut, is an indication that these dogs are killing more mouflon that they require to feed. Such remaining dead animals are becoming food for foxes, which are opportunistic and usually are after dead mouflon. When they locate one, they stay in the vicinity until nothing is left except the bones (Hadjisterkotis, 1992, 1996a; Constantinou & Hadjisterkotis, 2016).

To try to control the losses of sheep from predation, in 1958 the government assigned officials from the Forestry Department as full-time patrols armed with shotguns to kill dogs roaming unattended in the forest. In addition, meat injected with strychnine was used as a control agent until the late 1970’s. Over the period from 1985 to 1989 there were 15 reports of dogs in the forest. Of 22 dogs actually seen by foresters, seven (32%) were shot (Hadjisterkotis and Bider, 1992). Based on the above observations, Hadjisterkotis and Bider concluded that their limited data could support the hypothesis that dogs may exert some culling on diseased, parasitized or otherwise handicapped animals. The average number of dogs seen during those times was 1.5 dogs per group. Although full-time patrols armed with shotguns managed to kill about one third of the dogs seen in the forest, in the early 1990’s this practice was interrupted. Although in the 80’s there were one or two dogs each time, as I noted earlier, at least in the northern part of Pafos forest there are much larger groups of dogs.

Another factor that contributed to the reduction of dogs in the 1970’s and 1980’s was an antiechinococcosis campaign, which was implemented by the Department of Veterinary Services. Echinococcosis/hydatidosis was widespread in Cyprus before the 70s. It heavily infected almost every food animal and was a very serious public health problem. As a result of the seriousness of the disease, an antiechinococcosis campaign was implemented by the Department of Veterinary Services from 1971 until 1985. During this campaign over 85.000 stray dogs (the only final host of *Echinococcus granulosus* in Cyprus) were exterminated and about 14,000 bitches spayed. The cycle of the disease was considered completely interrupted, and infection in domestic sheep by 1985 was less than 0.1%, confined to very old animals. After 1985 it was believed and reported that echnococcosis/hydatosis had been eradicated from dogs, food animals and humans. During the last few years, there were only sporadic cases of hydatid cysts detected on slaughter. From November 1985 until January 1997, 47 Cyprian mouflon from Pafos forest were examined for Cystic echinococcosis. Only a 10 year old male was found infected. It had two fertile hydatid cysts on the lung, two and five cm in diameter (Hadjisterkotis, 1997). This is an indication that with the recent increase of dogs in the forest, the danger of increasing the cases of echinococcosis on the island is possible.

Although poaching and the observed increased dog predation is a direct or/and indirect result of small game hunting inside and around Pafos forest, the activities of hunters and their dogs predispose mouflon to a number of decimating factors, such as diseases, starvation, lack of nutrients, disruption of flock formation, etc. Limnitis is a valley with open grassy areas and many sharp cliffs which provide refuge for females for lambing during spring. In addition, the concentration of females attracts rams during the breeding season, which takes place at the end of October and during November. Because of the above, a large number of animals used to concentrate in this valley. Hunting, which takes place from the beginning of November until the end of December, coincides with the rutting season. By the end of the hunting season most of the females are pregnant and easier to get caught by dogs (Hadjisterkotis and Bider, 1992). During the rutting season the animals, particularly the males, are preoccupied with searching for estrus females and therefore are easier to get run over by a car on the road or to shoot, because they are easy to approach. By this time the feeding conditions inside the forest, are very poor. The long dry Mediterranean climate summers of Cyprus are hot and dry. Usually by June all grasses are dead, and forage is reduced in quantity and quality (Hadjisterkotis, 1993, 2016). As pasture forage matures, the protein content declines, fiber increases, and both forage intake and digestibility decline (NRC, 1985; Hadjisterkotis 1993, 2016). Most of the trees and shrubs inside Pafos forest contain antiherbivore substances, such as volatile oils, essential oils, tannic and gallic acids (Hadjisterkotis, 1993; Hadjisterkotis, 2001b). These oils inhibit activities in the rumen flora (Nagy and Tengerdy, 1967; Longhurts *et al*., 1968; Nagy and Tengerdy, 1968; Nagy and Regelin, 1977). Ninety-five percent of the diet of the Cyprus mouflon are grass and forbs which are free of toxins (Hadjisterkotis, 1996b; Hadjisterkotis, 2001). A seasonal chemical analysis of the major mouflon forage species from Pafos forest indicated that during late Summer and early Fall, grass, which is the main item in the diet of mouflon, was below the minimum requirements for maintenance required by sheep (Hadjisterkotis, 1993), which is 7% (NRC, 1985). The animals are further stressed at the end of October with the beginning of the rutting season, which lasts for about one month. Many males lose body weight during the rut, because the time that would be spent grazing is taken up with activities such as following estrous females and fighting rival males (Hadjisterkotis, 1993). The area of Limnitis valley is characterized by steep cliffs and high mountains. Because the energy cost of vertical travel is approximately 10 times the energy cost of horizontal travel (Clapperton 1964), animals living in this area need much more energy than the animals at the edge of the forest, and more energy to escape when disturbed. Hunter disturbance, at the edge of the forest, as well in Limnitis valley and around the villages Kampos and Tsakkistra during small game hunting as well as during the woodpigeon hunting seasons in August (Hadjisterkotis and Haaften, 1997: map 1, Hadjisterkotis, 1999), might force the animals to leave a good forage area and to seek refuge in more rocky parts of the forest, which decreases their condition and increases stress and therefore the mortality rate.

The low-quality food in combination with high loss of energy during the rut, the hunting activities that might disturb the animals during the rut, and the drop in winter temperature lead to a greater demand for energy. The above, possibly in combination with diseases and parasites, leads to a loss of condition and eventually death in both male and females. Therefore, as it was observed by Hadjisterkotis (2002), the highest mortality for Cyprus mouflon is during the rut and the period immediately following, for one or two months. Due to the rutting season, poor nutrition of the late summer and autumn, the drop in temperature and the above disturbances from dogs and hunters, the rams have no chance to recover and they die. Depending on the rains, fresh grass might not appear, particularly inside the forest until one and sometimes two months after the rut. Shortly after the rains the vegetation begins to grow, providing young, easily digestible material rich in protein and nutrients, especially forbs and grasses, which help the animals to recover. However, for many of these animals, because of the stress imposed on them from poachers, hunters, stray and hunting dogs, as well as the other factors noted earlier, they might succumb to starvations, diseases and parasites long before the start of the rains.

**References:**


Biddulph, J., (1884). On the wild sheep of Cyprus. Proc. Zool. Soc. London. 593-596.

Clapperton, J.L. (1964). The effect of walking upon the utilization of food by sheep. Br. J. Noutr. 18:39.

Garel, M., Marchand, P., Bourgoin, G., Santiago-Morerno, J., Portanier, E., Piegert, H., Hadjisterkotis, E., & Gugnasse, J-M., (2020). Mouflon Ovis gmelina Blyth, 1841. In K. Hackländer & F.E. Zachos (eds.) *Handbook of the Mammals of Europe* (pp. 1-35). Springer International Publishing. https://doi.org/10.10017/978-3-319-65038-8_34-1

Constantinou, G., & Hadjisterkotis, E. (2016). Fox predation on Cyprian mouflon and marine turtles on Cyprus. Pages 68-69, In: Hadjisterkotis E., (edr.) 6th World Congress on Mountain Ungulates and 5th International Symposium on Mouflon – Abstracts. August 29 – September 1st, Nicosia Cyprus. Ministry of the Interior, Nicosia.

Evripidou, P. (2010). Οργανωμένο κύκλωμα λαθροθηρίας - Υπάρχουν οικογένειες που συντηρούνται μονίμως από την αγορά του αγρινού. Αλήθεια, Δευτέρα 13th of September 2010.`

Guilaine, J., Briois, F., Coularou, J., & Carrére, I. (1995). L'établissement Néolithique de Shillourokambos (Parekklisha, Chypre) Premieres résultats. *Report of the Department of Antiquities, Cyprus:* 11-32.

Guilaine, J., & Briois, F. (2007). Shillourokambos and the neolithization of Cyprus: Some reflections. *Eurasian Prehistory* 4(1-2), 159–175.

Guilaine, J., Briois, F., & Vigne, J.-D. (eds.). (2011). *Shillourokambos. Un établissement néolithique pré-céramique à Chypre. Les fouilles du Secteur 1*. Editions Errance-Ecole Française d' Athènes, Paris. 1248 pp.

Haaften van J.L, (1974). European mouflon in the Mediterranean region. Zeitschrift fuer Jigdwissenschaft 20(4):181-184.

Hadjisterkotis, E. (2001). The Cyprus mouflon, a threatened species in a biodiversity "hotspot" area. Pages 71-81, In Nahlik A. and Walter Uloth (eds*.), Proceedings of the International Mouflon Symposium,* Sopron, Hungury.

Hadjisterkotis, E. (1992), The Cyprus mouflon *Ovis gmelini ophion*: Management, conservation and evolution. Ph.D. Thesis, McGill University, Montreal.

Hadjisterkotis E. (1996a). Fox and avian predation on Cyprus mouflon (*Ovis gmelina ophion*). Page 28 In: E. Hadjisterkotis (edr.) Abstracts – Second International Symposium on Mediterranean Mouflon. 17-20 April 1996, Nicosia, Cyprus. Game Fund, Ninistry of the Interior, Nicosia, Cyprus.

Hadjisterkotis, E. (1996b). Ernährungsgewonheiten des Zyprischen Mufflons *Ovis gmelini ophion* (Food habits of the Cyprus mouflon *Ovis gmelini ophion*) Z. Jagdwiss. 42 : 256-263.

Hadjisterkotis, E. (1997). The first record of *Echinococcus granulosus* in Cyprus mouflon *Ovis gmelini ophion*. Pages 84 in: Proceedings oft he 2nd World Conference on Mountain Ungulates, 5-7th May 1997, Saint-Viincent, Aosta, Italy.

Hadjisterkotis, E., (1999). Gefahren für das Zyprische aufgrund des Vorkommens als einzelne Restpopulation in einmem einzigen Verbreitungsgebiet (Dangers facing the Cyprus mouflon from being one population in one reserve). Z. Jagdwiss. 45:27-34.

Hadjisterkotis, E. (2000). Breeding Phenology and Success of the Woodpigeon (*Columba palumbus*) in Cyprus. Game and Wildlife Science, Vol. 17 (2): 81-92

Hadjisterkotis, E. (2002). Seasonal and monthly distribution of deaths of Cyprus mouflon *Ovis gmelini ophion*. Pirineos, 157: 81 a 88, JACA; 2002:81-88.

Hadjisterkotis, E. (2016). Population ecology and food habits off the Cyprus mouflon. Page 47. In: Hadjisterkotis, E. (edr). 6th world congress on Mountain Ungulates and 5th International Symposium on Mouflon – Abstractrs (3rd edn), August 29 – September 1st, 26, Nicosia Cyprus. Ministry of the Interior, Nicosia, Cyprus.

Hadjisterkotis E., & van Haaften, J.L. (1997). Die Niderwilldjagd im Wald von Paphos und ihre Auswirkungen auf gefahrdete zyprische Mufflon *Ovis gmelini ophion*. (Small game hunting in the forest of Paphos and its effects on the endangered Cyprian mouflon *Ovis gmelini ophion*) Z. Jagdwiss. 43: 279-282.

Hdjisterkotis, E., & Bider, J.R. (1993). Reproduction of Cyprus mouflon *Ovis gmelini ophion* in captivity and in the wild. Int. Zoo Yb. 32: 125-131

Hadjisterkotis E., & Bider, J.R. (1997). Chapter 6.4 Cyprus. In: D.M. Shackleton, *Wild Sheep and Goats and their Relatives, Status Survey and Conservation Action Plan for Caprinae*, IUCN/SSC Caprinae Specialist Group. Gland Switzerland.

Hadjisterkotis, E., & Lambrou, L. (2001). The role of the zoological garden of Limassol in wildlife conservation. Pages 103, In: E. Hadjisterkotis (edr.) XXVth International Congress of the International Union of Game Biologists I.U.G.B. and IXth International Symposium Perdix, Wildlife Management in the 21st Centary – Abstracts, September 3-7, 2001 Limassol, Cyprus.

Hadjisterkotis, E. (2001). The Cyprus mouflon, a threatened species in a biodiversity “hotspot” area. Pages 71-81, In Nahlik A. and Walter Uloth (eds.*), Proceedings of the International Mouflon Symposium,* Sopron, Hungury.

Hadjisterkotis, E. (2002). Seasonal and monthly distribution of deaths of Cyprus mouflon *Ovis gmelini ophion*. Pirineos, 157: 81 a 88, JACA; 2002:81-88.

Kassinis, N., Ioannou, I., Panayides, P., Mammides, C., Nicolaou, K. (2016). Current status, population dynamics and causes of mortality in Cyprus mouflon. Page 52, In: E. Hadjisterkotis (edr.) Proceedings of the 6th World Congress on Mountain Ungulates and 5th International Symposium on Mouflon – Abstracts, (3rd edn) August 29- September 1st, Nicosia Cyprus. Ministry of the Interior, Nicosia, Cyprus.

Longhurts, W.M., Oh, H.K., Jones, M.B., & keptner, R.E. (1968). A basis for palatability of deer forage plants. Trans. N. Amer. Wildl. And Nat. Res. Conf. 33:181-189.

Nagy, J.G. & Regelin, W.L. (1977). Influence of plant volatile oils on food selection by animals. Pages 225-230 In: Pterle, ed. XIII International congress of Game Biologists, The Wildlife Society, Wildlife Management Institute, Washington, D.C.1-538.

Nagy, J.G. & Tengerdy, R.P. (1967). Antibacterial action of essential oils of *Artemisia* as an ecological factor. I. Antibacterial action of the volatile oils of *Artemisia* *tridentanta* and *Artemisia nov*a on aerobic bacteria. Appl. Microbiol. 15(4):8129-821.

Nagy, J.G. & Tengerdy, R.P. (1968). Antibacterial action of essential oils of *Artemisia* as an ecological factor. I. Antibacterial action of the volatile oils of *Artemisia tridentanta* (big sagebrush) on bacteria from the rumen of mule deer. Appl. Microbiol. 16(4):441-4444..

National Research Council (1985). Nutrient Requirements of sheep. 6th edn. National Academy Press, Washington D.C. 1-99.

Nicolaou, H., Hadjisterkotis, E., & Papasavvas, K. (2016) (first edition). Past and present distribution and abundance of the Cyprus mouflon. Page 103, In: E. Hadjisterkotis (edr.) Proceedings of the 6th World Congress on Mountain Ungulates and 5th International Symposium on Mouflon – Abstracts (3rd edn), August 29- September 1st, Nicosia Cyprus. Ministry of the Interior, Nicosia, Cyprus.

Toumazos, P., & Hadjisterkotis, E. (1997) Diseases of the Cyprus mouflon as determined by Standard gross and histopathological methods. Pages 150-161 *in* E. Hadjisterkotis (ed.). *Proceedings of the Second International Symposium on Mediterranean Mouflon*. Game Fund, Nicosia, Cyprus.

Tsintides, T., Kakouris, E., Hadjisterkotis, E. (2016). (1st edn) Pafos forest, a biodiversity “hospot” and center of endemism: measures for preventing forest fires fro the conservation and management of mouflon. Page, 91. In: E. Hadjisterkotis (edr.). Proceedings of the 6th World Congress on Mountain Ungulates and 5th International Symposium on Mouflon – Abstracts, August 29- September 1st, Nicosia Cyprus. Ministry of the Interior, Nicosia, Cyprus.

Vigne, J.-D., Briois, F., Zazzo, A., Willcox, G., Cucchi, T., Thiébault, S., Carrère, I., Franel, Y., Touquet, R., Martin, C., Moreau, C., Comby, C. & Guilaine, J. (2012). The first wave of cultivators spread to Cyprus earlier than 10,600 years ago. *Proceedings of the National Academy of Sciences USA* 109(22): 8445–8449.

Vigne, J.-D., Zazzo, A., Saliège, J.-F., Poplin, F., Guilaine, J. & Simmons, A. (2009). Pre-Neolithic wild boar management and introduction to Cyprus more than 11,400 years ago. *Proceedings of the National Academy of Sciences USA* 106(38): 16131–16138.

Vigne, J.-D., Zazzo, A., Cucchi, T., Carrère, I., Briois, F. & Guilaine J. (2014). The transportation of mammals to Cyprus sheds light on early voyaging and boats in the Mediterranean sea**.** *Eurasian Prehistory* **10** (1–2): 157–176.
